# Improving the heterologous production of fungal peroxygenases through an episomal *Pichia pastoris* promoter and signal peptide shuffling system

**DOI:** 10.1101/2020.12.23.424034

**Authors:** Pascal Püllmann, Martin J. Weissenborn

## Abstract

Fungal Peroxygenases (UPOs) have emerged as oxyfunctionalization catalysts of tremendous interest in recent years. However, their widespread use in the field of biocatalysis is still hampered by their challenging heterologous production, substantially limiting the panel of accessible enzymes for investigation and enzyme engineering. Building upon previous work on UPO production in yeast, we have developed a combined promoter and -signal peptide shuffling system for episomal high throughput UPO production in the industrially relevant, methylotrophic yeast *Pichia pastoris*. 11 endogenous and orthologous promoters were shuffled with a diverse set of 17 signal peptides. Three previously described UPOs were selected as first test set, leading to the identification of beneficial promoter/signal peptide combinations for protein production. We applied the system then successfully to produce two novel UPOs: *Mfe*UPO from *Myceliophthora fergusii* and *Mhi*UPO from *Myceliophthora hinnulea.* To demonstrate the feasibility of the developed system to other enzyme classes, it was applied for the industrially relevant lipase CalB and the laccase Mrl2. In total, approximately 3200 transformants of eight diverse enzymes were screened and the best promoter/signal peptide combinations studied at various co-feeding, derepression and induction conditions. High volumetric production titers were achieved by subsequent creation of stable integration lines and harnessing orthologous promoters from *Hansenula polymorpha*. In most cases promising yields were also achieved without the addition of methanol under derepressed conditions. To foster the use of the episomal high throughput promoter/signal peptide *Pichia pastoris* system, we made all plasmids available through Addgene.

## INTRODUCTION

Fungal unspecific peroxygenases (UPOs) have gained substantial interest as versatile oxyfunctionalization biocatalysts since their initial description in 2004.^1^ UPOs are solely relying on hydrogen peroxide as co-substrate for the formation of the reactive oxyferryl (Compound I) intermediate. In contrast to the well-known cytochrome P450 enzymes, they do not rely on auxiliary electron transport proteins and expensive cofactors.^2–3^ Recent studies demonstrate impressive activities on a wide variety of structurally diverse substrates.^4–6^

Arguably the major bottleneck in the widespread use of UPOs, naturally occurring as disulphide-bridged, glycosylated, secreted enzymes remains their challenging heterologous production.^2, 7–8^ Recent work describes the production of three UPOs in *E. coli,* however the reported recombinant yields are comparably low and proof for the high throughput capacity of the developed prokaryotic system is lacking.^9–10^ Utilizing the eukaryotic host *Saccharomyces cerevisiae, AaeUPO* derived from *Agrocybe aegerita* was engineered for active secretion to a titre of 8 mg/L.^11^ Subsequent directed evolution approaches based on this yeast-secretion variant AaeUPO* led to improved variants for the synthesis of 1-naphthol and 5’-hydroxypropranolol.^12–13^The UPO secretion variant could also be successfully produced utilizing the methylotrophic yeast *Pichia pastoris* (syn. *Komagataella phaffii*).^14^

*P. pastoris* represents an industrially highly relevant heterologous production platform due to beneficial metabolic characteristics as well as a steadily growing toolbox of valuable synthetic biology parts such as modular circuits, strong as well as tightly regulated promoters and signal peptides.^15–17^ Especially promoter elements are popular, which are derived from genes involved in the specialized methanol utilization (MUT) pathway. The most famous and commonly used MUT based promoter, AOX1 is derived from the alcohol oxidase 1 gene. The widespread application of these promoters is based on their common tight repression and strong methanol induction profile.^18–19^ Besides this sharp repression vs. induction profile, several MUT promoters were described in recent years, enabling methanol-free derepressed gene expression upon carbon source depletion. This derepressed gene expression is of high interest as methanol bears drawbacks as inducer in the course of large-scale fermentations as toxic and flammable agent.^19–20^ Recent studies report orthologous MUT promoter elements derived from the methylotrophic yeast *Hansenula polymorpha* (syn. *Ogataea angusta*) outperforming endogenous, strong *P. pastoris* MUT promoters for target protein production.^21–22^

Despite being a heterologous host of outstanding interest, the use of *P. pastoris* for high throughput enzyme engineering endeavours remains rather limited to few examples.^23–27^ This is likely due to a limited number of suitable autonomously replicating sequences (ARS) for P. *pastoris,* which confer episomal stability. For the model yeast organism *S. cerevisiae,* a multitude of directed evolution examples exist utilizing well-known episomal systems.^11^, ^28–29^

In this work we present a high throughput episomal expression system for *P. pastoris*. The system is based on principles of modular Golden Gate cloning^30–32^ and enables the rapid assessment of the suitability of promoter/signal peptide combinations for recombinant protein secretion. 11 orthologous or endogenous MUT promotors can be combined with 17 signal peptides for each individual gene of interest, leading to 187 unique combinations. This system was validated and optimized using known UPOs and further enabled the first yeast production of *DcaUPO* and the discovery of two new UPOs. To demonstrate versatility of the system, ideal episomal combinations were additionally determined for the lipase CalB and the laccase Mrl2. After the screening of approx. 3200 primary transformants in the episomal *P. pastoris* setup, the best performing transcription units were genomically integrated and led to high production titres within shake flask production setups.

## RESULTS

### Building and testing the setup

We selected eleven MUT promoters: eight derived from *P. pastoris (P_P_p_PMP20_; P_P_p_FLD1_; P_P_p_FDH1_; P_P_p_DAs1_, P_P_p_DAS2_, P_P_p_CAT1_; P_P_p_Aox1_, P_P_p_ALD4_)* and three orthologous promoters from *H. polymorpha (P_H_p_MOX_; P_H_p_DHAS_; P_H_p_FMD_*).^17, 19, 21^ All promoters were cloned as Level 0 modules into pAGM9121 bearing an identical Kozak sequence (^-10^ATCATACAAA^0^**ATG**). The design was implemented to be fully compatible with the previously designed yeast secretion system (Supplemental Figure 1).^33^ A list of all promoter and signal peptide sequences is given (Supplemental Tables 2 and 3). Tetrapartite expression units (**5’** promoter - signal peptide - gene - C-terminal Tag **3’**) are assembled in a one-pot, one-step Golden Gate reaction within the episomal yeast expression plasmid pPAP004, which can be propagated in *E. coli* as well as *P. pastoris* (Figure 1 A). Identified beneficial promoter-signal peptide combinations can be swapped as complete transcription unit into an integrative plasmid (pPAP003) in the course of a second Golden Gate reaction leading to stable strains for high yield and antibiotic-free protein production. As a first test object, the recently described enzyme *Mth*UPO was chosen.

**Figure 1.**
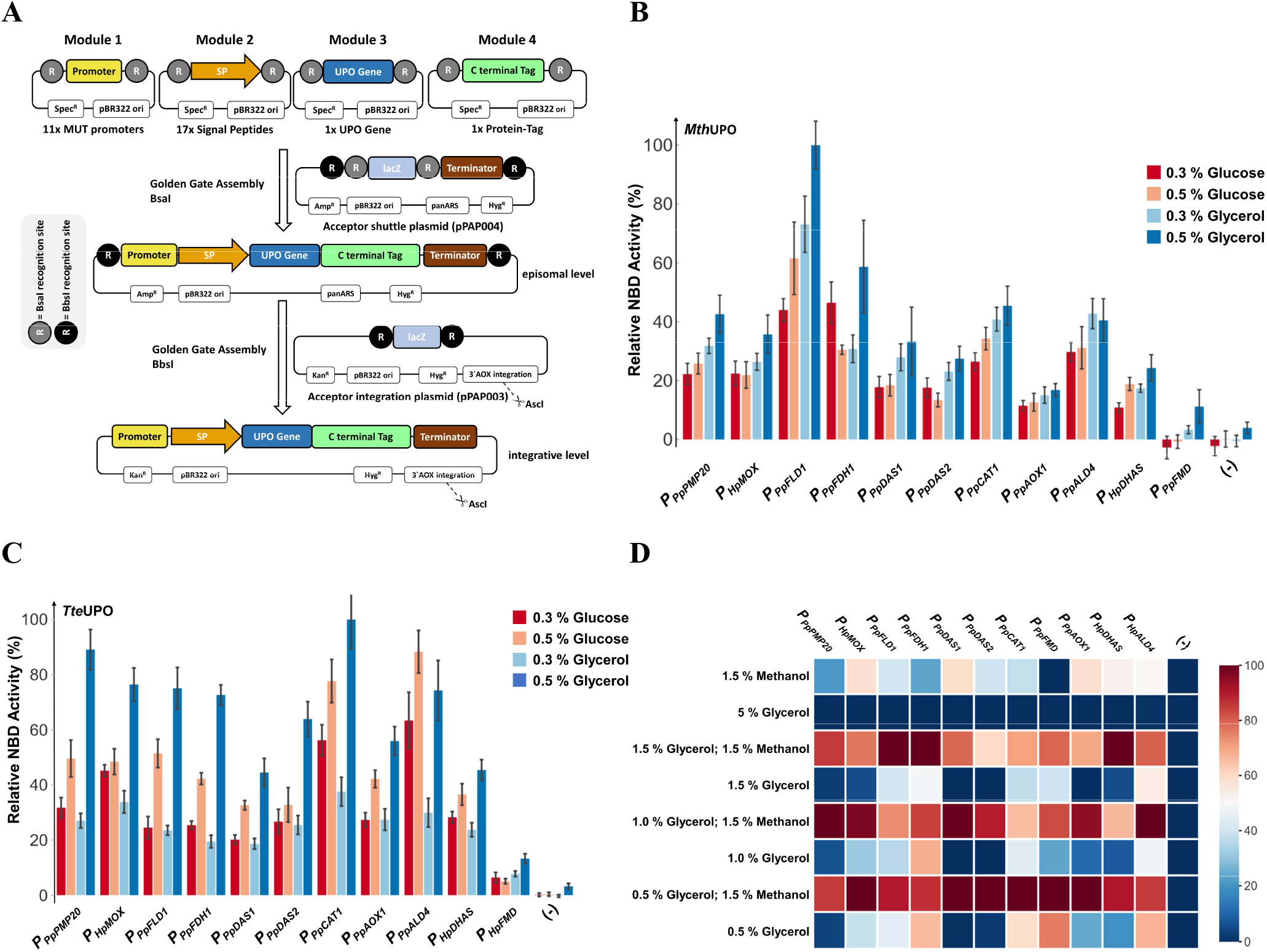
Design and testing of the episomal MUT expression shuffling system in *P. pastoris.* A: Episomal tetrapartite expression units (5’to 3’: MUT promoter-signal peptide-UPO Gene-C-terminal tag) can be assembled within an *E. coli/P. pastoris* shuttle plasmid (pPAP004) and transformed yeast clones are screened for activity. In a second Golden Gate reaction, the previously identified, suitable expression units can be transferred to another plasmid (pPAP003) for stable genomic integration, yielding strains for high yield protein production. B: Testing of individual MUT promoter strengths using MthUPO as test enzyme (signal peptide: *“Sce-*α Galactosidase”). 6 biological replicates were cultivated within the MTP setup and UPO specific NBD conversion were measured in the supernatant after 72 h cultivation. (-) in every case indicates the empty plasmid control (pPAP004). The highest mean activity was set as 100 % and all other means were normalized relative to this value, standard deviations are indicated. Carbon sources were used in a co-feeding system utilizing two sources, glucose or glycerol as indicated and 1.5 % methanol in all cases. C: Analogous setup to B but using TteUPO as test enzyme (signal peptide: *“Sce-* Invertase 2”). D: Carbon source screening of TteUPO constructs. Respective MUT constructs as before (similar to setup C) tested under 8 different carbon source conditions utilizing methanol induced (0.5 % to 1.5 % glycerol plus 1.5 % methanol and 1.5 % Methanol) and methanol-free conditions (0.5 to 5 % glycerol). Cultivation, screening and analysis as described before. The highest mean activity of every promoter was set to 100 % and all other mean values of the respective promoter normalized accordingly leading to the heatmap as shown. An absolute UPO activity chart is depicted in Supplemental Figure 2 based on the same activity measurements.

In the initial approach, all MUT promoter constructs were built in combination with the previously identified suitable signal peptide “*Sce*-α Galactosidase” for the secretion of MthUPO derived from *Myceliophthora thermophila.^33^* This approach led to 11 defined constructs. *P. pastoris* cells were cultivated for 72 h in 96 well plates using a co-feeding strategy of the primary carbon source glucose or glycerol (0.3 or 0.5 % (v/v)) and the inducing carbon source (1.5 % (v/v) methanol). Except for *P_H_p_FMD_*, all promoters led to a distinguishable activity using the UPO specific NBD Assay^34^ and when compared to the negative control (pPAP004) under all tested co-feeding conditions (Figure 1 B). In general, *P_P_p_FLD1_* appeared to be the most suitable promoter for MthUPO production, and the co-feeding conditions using 0.5 % glycerol led to the highest average activity of all promoters.

As a second test, the previously described enzyme TteUPO from *Thielavia terrestris* was selected (Figure 1 C). TteUPO was combined with all respective promoters following the same principles as before but combined with the signal peptide “*Sce*- Invertase 2”.^33^ General observations were similar to the screening of MthUPO, but identifying *P_P_p_CAT1_* as most suitable promoter within this setup and obtaining clear UPO activities for all promoters including *P_H_p_FMD_.*

Since some of the included promoters were previously described to gain activity under conditions of primary carbon source depletion (derepression) we further wanted to test the UPO production behaviour of the 11 MUT promoter under varying derepressive/inductive carbon sources.^18–19, 21, 35^ We chose various co-feeding (0.5 to 1.5 % glycerol + 1.5 % methanol), derepressive (0.5 to 1.5 % glycerol), excess (5 % glycerol) and solely methanol (1.5 %) conditions. The negative control (pPAP004) and all MUT samples under excess glycerol conditions did not exhibit any UPO activity (Figure 1 D). This tight repression demonstrated that promoter activity and UPO production are solely based on methanol induction or carbon source depletion based derepression. In all cases, the highest UPO production rates were obtained employing a co-feeding strategy using 0.5 % (5 promoters) or 1 to 1.5 % glycerol (3 promoters each). Interestingly, using methanol as sole carbon source and inducing agent, resulted in a reduced cell density and decreased UPO activity (approx. 20 - 60 % of maximal activity). This observation highlights the appeal of the co-feeding setup for optimal protein production. Based on the observed production patterns within this setup, the MUT promoters can be grouped in solely methanol inducible *(P_P_p_PMP20_; P_P_p_DAS1_; P_P_p_DAS2_*), slightly derepressed *(P_H_p_MOX;_ P_H_p_DHAS_; P_P_p_AOX1_),* medium *(P_P_p_FLD1_)* and strongly derepressed *(P_P_p_FDH1_; P_P_p_CAT1_; P_P_p_ALD4_; P_H_p_FMD_).* Time course investigations were performed from 20 to 100 h after inoculation of *Tte*UPO constructs utilizing six derepressed promoters (Supplemental Figure 3). In general, it could be observed that UPO activity primarily accumulates in the supernatant in the interval between 30 and 50 h, correlating with reaching the maximal cell density within the wells. After that, the UPO activity slightly increased until the endpoint (100 h) within most setups. Interestingly, testing range spanning derepressed conditions (0.5; 1.0 or 1.5 % glycerol) resulted in no substantial differences in the activity pattern of any promoter. We did not obtain any evidence for an occurring lag-phase of derepressed UPO production when using higher glycerol concentrations (1.5 %) over low concentrations (0.5 %).

### Combining signal peptide and promoter shuffling

In previous studies using two promoters and various signal peptides, we observed rather surprising effects of certain combinations on the UPO yield.^33^ The classical model of selecting a strong promoter and a suitable signal peptide for secretion was valid in many cases. Still, individual results pointed towards pronounced synergistic effects of chosen promoter/signal peptide combinations on the UPO activity rather than selecting a strong promoter to drive target gene expression. We built and tested 18 TteUPO constructs to further investigate this preliminary observation. Three previously as suitable identified promoters *(P_H_p_MOX_; P_p_p_ALD4_; P_P_p_CAT1_)* and six suitable signal peptides were employed (Supplemental Figure 4). In three cases with the signal peptides “*Sce*-Prepro”, “*Sce*-Invertase 2” and “*Sce*-Killer Protein” an expected promoter-based pattern of UPO activity was observed: *P_P_p_CAT1_ > P_H_p_MOX_* ≥ *P_P_p_ALD4_.* In two cases (“Mro-UPO” and “Kma-Inulinase”), the obtained UPO activity of the *P_P_p_CAT1_* construct was approx. doubled in comparison to the other promoters. Even more surprising, this observation was inverted in one case (“Hsa-Serum Albumin”) towards *P_H_p_MOX_* outperforming the other two promoters by twofold. These results further strengthen a more synergistic hypothesis of episomal protein secretion in *P. pastoris*, rather than just taking promoter strength and signal peptide suitability into consideration as isolated factors.

Based on this synergism hypothesis, we performed all subsequent enzyme screenings as combined promoter-signal peptide shuffling approaches with 11 MUT promoters and 17 signal peptides leading to 187 unique combinations. The resulting libraries were transformed into *P. pastoris* and 370 to 378 transformants of each library screened for UPO specific NBD activity using a co-feeding of 0.5 % glycerol and 1.5 % methanol (Figure 1 D). Initial screening of the libraries of TteUPO, MthUPO, and AaeUPO*^11,14^ revealed differing patterns. *Tte*UPO and *Mth*UPO exhibited a high promiscuity regarding functional promoter/signal peptide combinations. 66 % (TteUPO) and 56 % (MthUPO) of all transformants exhibited UPO activity (Supplemental Figure 5 and 6). For *Aae*UPO*, however, just a small fraction of 7 % of the transformants were active (Supplemental Figure 7). These observations are consistent with previous findings performing a signal peptide shuffling approach with these enzymes.^33^ Among the top 15 most active hits of each enzyme, a vast diversity of shuffled promoters (TteUPO: 5; MthUPO: 6; AaeUPO*: 7) and signal peptides parts (TteUPO: 8; MthUPO: 8; AaeUPO*:2) could be observed (Figure 2 A). The low diversity of signal peptides for AaeUPO* is in good agreement with previous findings that suggested low promiscuity of this enzyme towards the signal peptide panel.^33^ Subsequent carbon source dependent activity screening was performed of the top two performing episomal (abbreviated with (e) and their integrative counterparts (i) (Figure 2 B). These carbon source variations revealed a diverse dynamic of UPO production under derepressed and induced conditions. The *H. polymorpha* derived promoter *P_H_p_FMD_* proved to be the best performing promoter in case of all enzymes. The highest NBD activity under episomal production however varied for every UPO. The highest levels were obtained under co-feeding conditions for TteUPO (1.0 % glycerol; 1.5 % methanol) and MthUPO (0.5 % glycerol; 1.5 % methanol) and for AaeUPO* even under methanol-free conditions with 1.0 % glycerol. Therefore, a “sweet spot” for every episomal construct can be found by carbon source screening. The ideal carbon source condition can then be further exploited as a standard condition to obtain maximal enzyme yields easing high throughput directed evolution endeavors. Utilizing one step TwinStrep-based affinity purification, all UPOs were obtained in high purity and analyzed for their extent of glycosylation and molecular weight by SDS PAGE, obtaining apparent molecular weights of 27 kDa (TteUPO), 35 kDa (MthUPO) and 45 kDa *(AaeUPO*,* Figure 2 C). Interestingly, when producing *Tte*UPO before in the identical *P. pastoris* strain but utilizing a different signal peptide (“*Sce*-Prepro”), we obtained a deglycosylated apparent MW of 35 kDa.^33^ This observation and the calculated molecular weight based on the enzyme’s primary sequence of 32 kDa suggests a occurring divergent cleavage of an N-terminal part of the mature enzyme.

**Figure 2.**
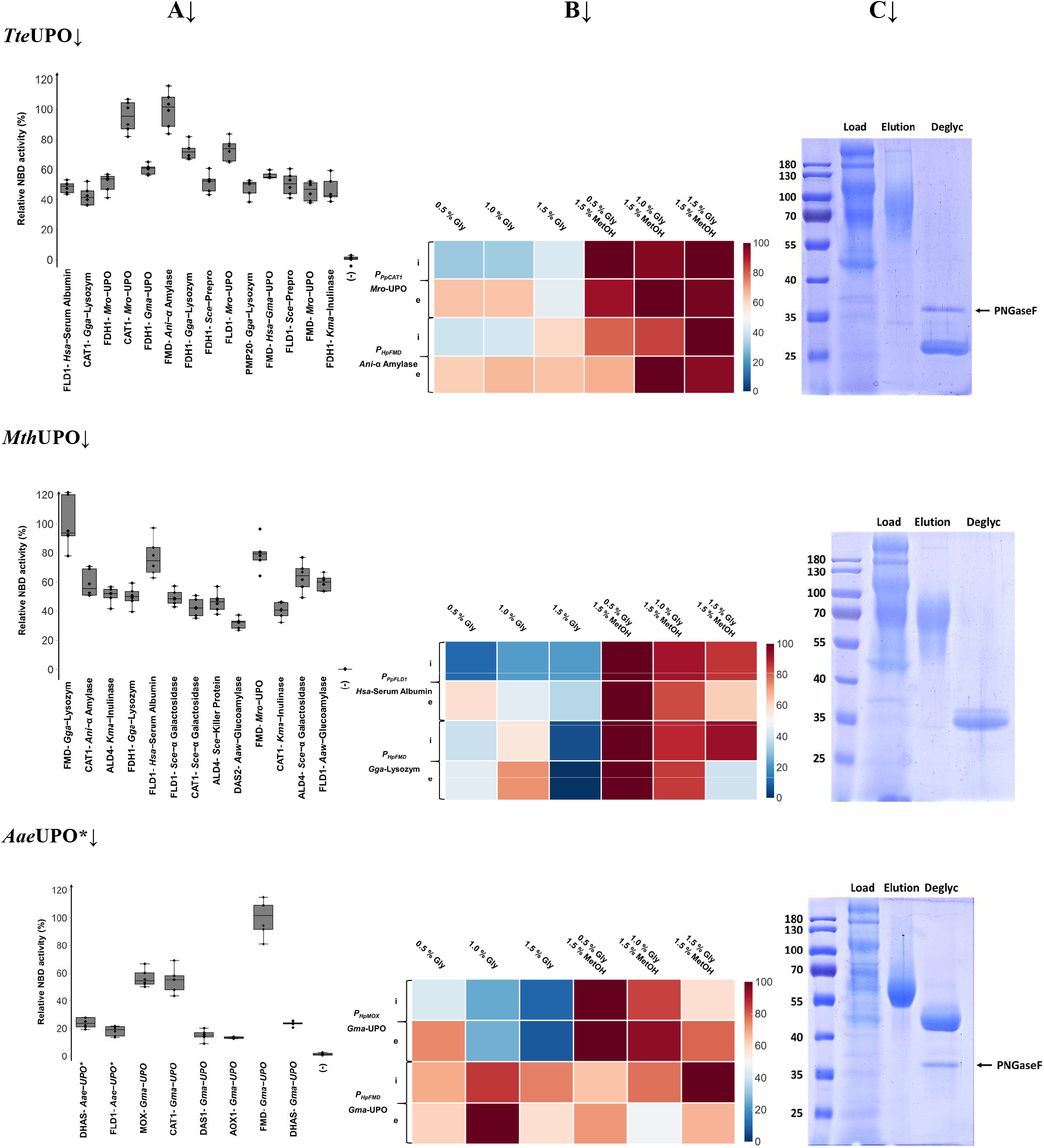
Simultaneous MUT promoter and signal peptide shuffling to improve heterologous UPO production. Three previously described UPOs (TteUPO, *MthUPO,* AaeUPO*) were subjected to one-pot, one-step promoter/signal peptide shuffling (187 possible unique combinations). A: Relative activity of non-redundant constructs among the most active 15 clones. 6 biological replicates were cultivated within the MTP setup and UPO-specific NBD conversion measured in the supernatant after 72 h cultivation. (-) in every case indicates the empty plasmid control (pPAP004). The highest mean activity was set as 100 % and all other data points were normalized relative to this mean value. B: Comparison of top two episomal (e) and integrative constructs (i) regarding UPO activity utilizing differing carbon source production conditions (screening conditions as described in A). The highest mean activity of every group was set to 100 % and all values normalized accordingly. Primary data are displayed in Supplemental figure 8. C: One step TwinStrep-based purification of recombinant UPOs. Load (50 ml sample after ultrafiltration), pooled elution (Elution) and elution fraction after enzymatic N-deglycosylation (PNGaseF treatment; Deglyc) were analyzed by SDS-PAGE (12 %).

### Introducing novel UPOs into the modular system

Taking the recently characterized *Mth*UPO as search query, we retrieved the sequences of two close homologs from a recent patent.^36^ The UPO from *Myceliophthora fergusii* (*Mfe*UPO) and from *Myceliophthora hinnulea* (*Mhi*UPO) share a high sequence identity of 91 % and 96 % towards MthUPO. As the third UPO, we selected the *DcaUPO* derived from *Daldinia caldariorum,* recently produced in *E. coli.*^10^ DcaUPO has a moderate sequence identity of 53 % and 52 % towards MthUPO and *Tte*UPO, respectively. The three new enzymes were subjected to the promoter/signal peptide shuffling approach. The resulting libraries were screened for UPO-specific activity using colorimetric screening assays with NBD or DMP as substrate. Primary screening landscapes revealed comparable proportions of active transformants for *Dca*UPO (26 %), *Mfe*UPO (24 %), and MhiUPO (31 %, Supplemental Figures 9 to 11). These proportions point towards a less pronounced promoter/signal peptide promiscuity than MthUPO (56 %) and TteUPO (66 %), but higher than *Aae*UPO* (7 %). Since the signal intensity of the NBD assay in case of the enzymes appeared to be rather low, further screenings of the three novel UPOs were conducted employing the more sensitive DMP assay.^12^

The top 15 hits exhibited a redundant pattern, due to the performed oversampling, limiting the number of unique constructs to a range between 6 (*Mfe*UPO) and 10 (*Dca*UPO, Figure 3 A). Nevertheless, as before a diverse panel of promoters (DcaUPO: 7; MfeUPO: 5; MhiUPO: 5) and signal peptides (DcaUPO: 6; *Mfe*UPO: 4; *Mhi*UPO: 7) was observed. Carbon source activity screening identified two *H. polymorpha* promoters occurring within the best performing constructs: *P_H_p_FMD_* (MfeUPO) and *P_H_p_MOX_* (*Mhi*UPO, *Dca*UPO, Figure 3 B). As shown before for all enzymes, a specific “sweet spot” of maximal activity within the episomal system was identified, while in case of *Dca*UPO co-feeding (0.5 % glycerol; 1.5 % methanol) led to maximal activity, *Mfe*UPO and *Mhi*UPO activity reached its maximum under derepressed condition with 1 % glycerol.

**Figure 3.**
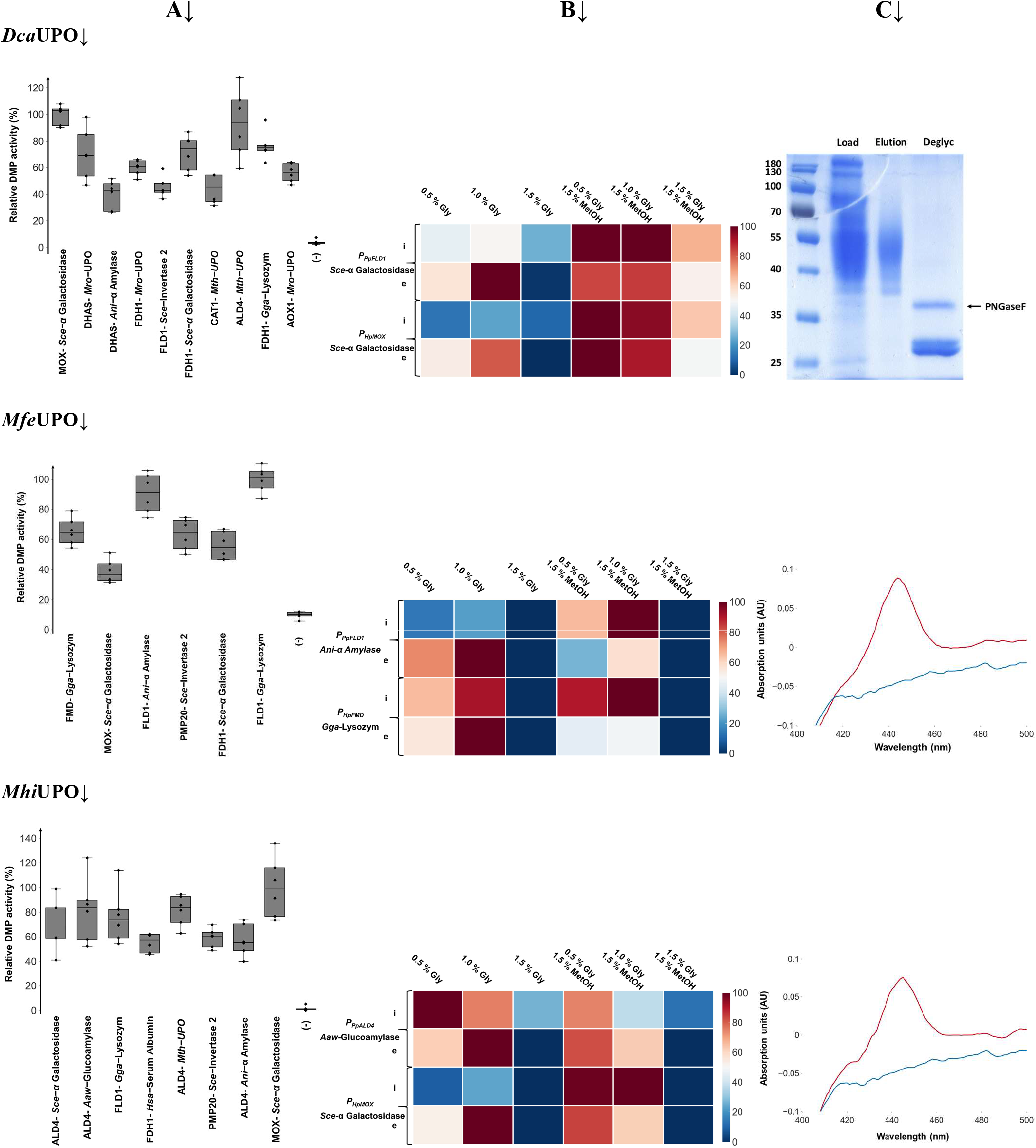
Testing the heterologous production of novel UPOs. Two previously UPOs (MhiUPO and *MfeUPO)* and a recently described UPO (DcaUPO) were subjected to one-pot, one-step promoter/signal peptide shuffling (187 possible unique combinations). A: Relative activity of non-redundant constructs among the most active 15 clones. 6 biological replicates were cultivated within the MTP setup and UPO-specific DMP conversion measured in the supernatant after 72 h cultivation. (-) in every case indicates the empty plasmid control (pPAP004). The highest mean activity was set as 100 % and all other data points normalized relative to this mean value. B: Comparison of top two episomal (e) and integrative constructs (i) regarding UPO activity utilizing differing carbon source production conditions (screening conditions as described in A). The highest mean activity of every group was set to 100 % and all values normalized accordingly. Primary data are displayed in Supplemental Figure 12. C: One step HIC-based purification of recombinant DcaUPO. Load (50 ml sample after ultrafiltration), pooled elution (Elution) and elution fraction after enzymatic N-deglycosylation (PNGaseF treatment; Deglyc) were analyzed by SDS-PAGE (12 %). *MfeUPO* and MhiUPO (red) were analyzed after ultrafiltration by differential CO spectra in comparison with the integrative negative control (pPAP003;

Purification of the three UPOs proved to be challenging. They could not be purified by means of affinity based TwinStrep purification. This is most likely caused by masking of the C-terminal TwinStrep tag due to pronounced glycosylation or might point towards proteolytic cleavage of the tag. Using classical purification techniques like ion exchange (IEX) and hydrophobic interaction chromatography (HIC), the UPOs could be captured, but specific elution was not achieved. UPO activity was rather spread over all elution fractions (salt gradient) pointing towards extensive glycosylation. This heavy glycosylation leads to a very heterogeneous enzyme pool with divergent physicochemical behavior, which are underlying principles of IEX and HI chromatography. Using OctylSepharose as HIC material, we obtained a tightly bound, pure fraction of *Dca*UPO by elution with ethylene glycol. SDS PAGE analysis (Figure 1 C and Supplemental Figure 13) revealed high purity as well as an extensive glycosylation with an apparent molecular weight of 40 to 55 kDa, which was reduced to approx. 27 kDa after deglycosylation. This apparent molecular weight reduction is in good agreement with the calculated molecular weight of *Dca*UPO. In the case of *Mfe*UPO and *Mhi*UPO, purification was not successful, but precise heme CO differential spectra were obtained compared with the integrative negative control pPAP003 (Figure 3 C and Supplemental Figure 14).

### Expanding the system to other enzyme classes

After successfully applying the episomal system for six UPOs we wanted to further expand the system towards other enzymes with a general challenging expression. We chose the widely used lipase B from *Candida antartica* (CalB) and a laccase derived from the basidiomycete *Moniliophthora roreri* (Mrl2).^37^ Both enzymes were subjected to promoter/signal peptide shuffling. Primary screenings were conducted using well-established colorimetric assays: 4-Nitrophenyl laurate for CalB and ABTS for Mrl2. The screening landscaping revealed medium to high promiscuity for CalB (Supplemental Figure 15; 44 %) and the overall highest rate of active transformants in the case of Mrl2 (70 %). Regarding Mrl2 (Supplemental Figure 16), we observed a pyramid-like shape of the activity landscape when ranking the clones based on activity. This shape differed substantially from the negative-exponential (very few clones with high activity, many medium-high activity clones) shaped landscapes for the other seven enzymes. Assessing the 15 most active transformants revealed again a grand diversity of occurring promoters (CalB: 4; Mrl2: 4) and signal peptides (CalB: 7; Mrl2: 9, Figure 4 A). Subsequent carbon source activity profiling of the top two constructs (Figure 4 C) confirmed *P_H_p_FMD_* as the most suitable promoter. With this promoter, the highest lipase/laccase activities were reached within the episomal system when a co-feeding was applied: CalB with 1.0 % glycerol and for Mrl2 with 0.5 % glycerol.

**Figure 4.**
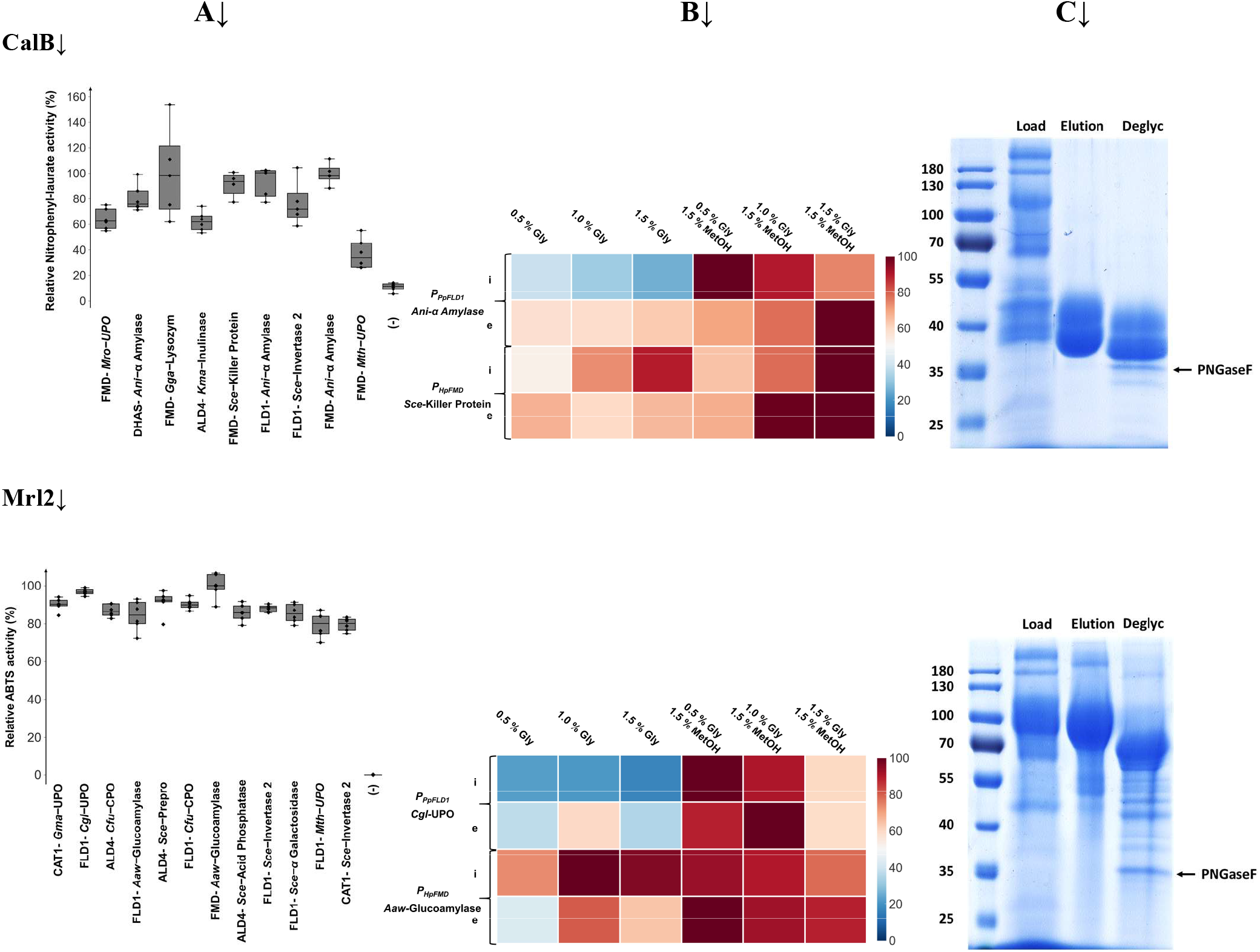
Expanding the dual shuffling system to other enzyme classes. Lipase CalB and laccase Mrl2 were subjected to one-pot, one-step promoter/signal peptide shuffling (187 possible unique combinations). A: Relative activity of non-redundant constructs among the most active 15 clones. 6 biological replicates were cultivated within the MTP setup and lipase specific (4-Nitrophenyl laurate) and laccase specific (ABTS) conversion measured in the supernatant after 72 h cultivation. (-) in every case indicates the empty plasmid control (pPAP004). The highest mean activity was set to 100 % and all other data points were normalized relative to this mean value. B: Comparison of top two episomal (e) and integrative constructs (i) regarding lipase (4-Nitrophenyl laurate) or laccase (ABTS) activity utilizing differing carbon source production conditions (screening conditions as described in A). The highest mean activity of every group was set to 100 % and all values normalized accordingly. Primary data are displayed in Supplemental figure 17 C: One step TwinStrep-based purification of recombinant enzymes. Load (50 ml sample after ultrafiltration), pooled elution (Elution) and elution fraction after enzymatic N-deglycosylation (PNGaseF treatment; Deglyc) were analyzed by SDS-PAGE (12 %).

Contrary to the difficult purification of the novel UPOs, CalB and Mrl2 were purified successfully employing TwinStrep affinity chromatography (Figure 4 C). The obtained apparent molecular weights for the glycosylated (CalB: 40 kDa; Mrl2: 100 kDa) and deglycosylated form after PNGaseF treatment (CalB: 38 kDa; Mrl2: 70 kDa) are in good agreement with reported _data._^37–38^

### Volumetric production yields

Besides the primary focus on the development of a modular episomal system, we also implemented an integrative plasmid (pPAP003; Figure 1 A) as second layer of the system. The integrative plasmid enables the rapid construction of stable strains for large scale, antibiotic-free protein production. Selecting the best performing, episomal constructs of all eight previous enzymes, we performed genome integration and subsequent shake flask cultivations in a 1-liter format. All enzymes were cultivated and produced without any further production optimization for 72 h (25 °C) under derepressed (1 % glycerol as sole carbon source) and induced conditions (start on 0.4 % glycerol; 1 % methanol spikes after 24 and 48 h). The volumetric yields of all enzymes under both conditions are displayed in Table 1. Especially when using *P_H_p_FMD_*, already under derepressed conditions, moderate (*Mth*UPO: 17 %) to high yields (CalB: 72 %) were obtained compared to the maximum yields with methanol induction. The promoter *P_H_p_-MOX_* proved to be relatively inactive under the tested derepression conditions, leading to no (*Dca*UPO) or marginal (MhiUPO) protein production. The novel enzymes *MfeUPO* and MhiUPO were produced with a titer of 6.5 and 5.7 mg/L respectively. We were able to surpass previously reported heterologous yields of TteUPO (+ 56 %) and AaeUPO* (+ 58 %).^11, 33^ An even more pronounced increase was achieved for *Dca*UPO with a 482 % improvement relative to the previous reports on production in *E. coli.^10^* Heterologous production of *Mth*UPO was not improved further but was achieved on a similar high level (22.4 mg/L).^33^ The volumetric yield of the laccase Mrl2 was increased by 191 % reaching the overall highest volumetric yield of 30.9 mg/L observed in the whole test setup. Solely the production level of CalB could not compete with previously published work.^38^

**Table 1.**
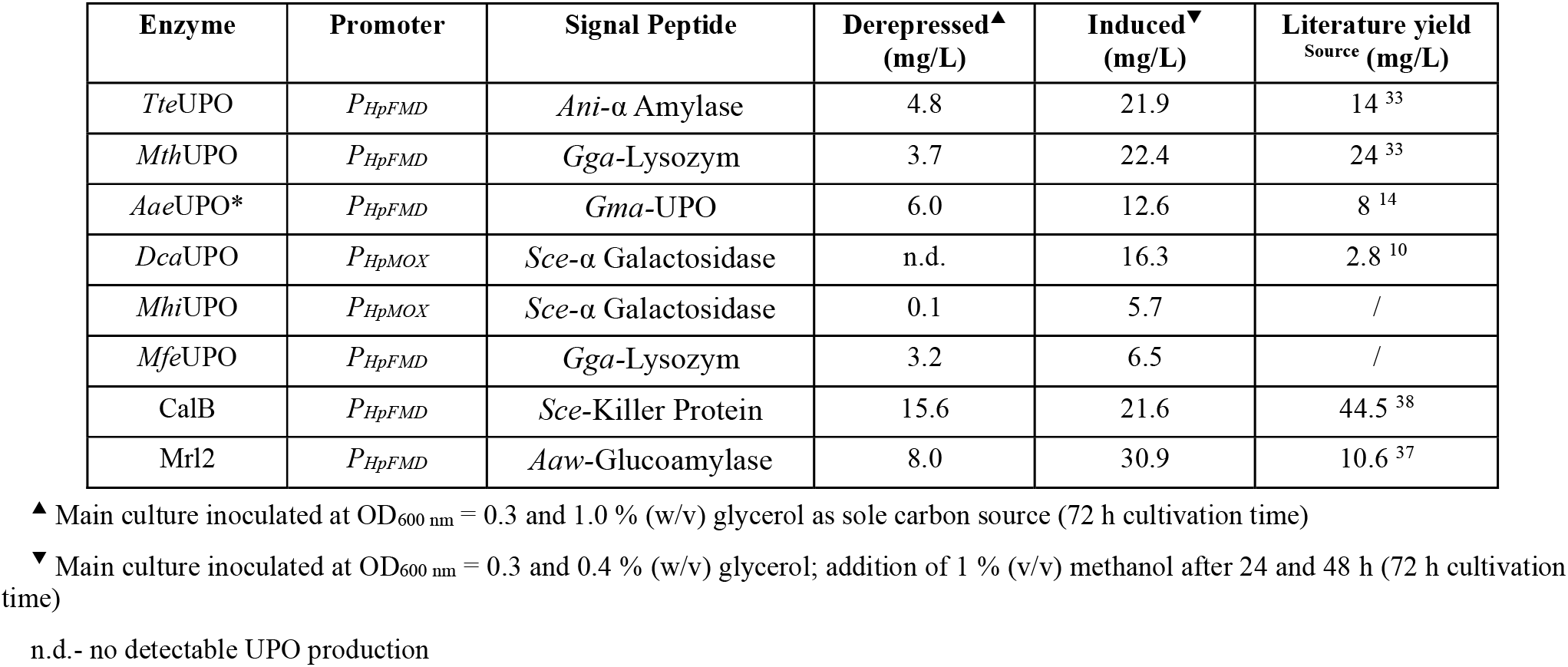
Volumetric Protein yields of recombinant enzymes

### Expanding the system to *H. polymorpha*

In the first report of the utilized ARS sequence (panARS), the authors also describe episomal stability in the yeast *H. polymorpha*.^39^ *H. polymorpha* is, analogous to *P. pastoris,* a methylotrophic yeast with outstanding interest in industrial biotechnology.^40^ We were intrigued, if our constructed episomal plasmid series based on the pPAP004 backbone also functions in *H. polymorpha*, conferring episomal stability and enabling target protein production. We transformed *H. polymorpha* with the *Tte*UPO promoter library (Figure 1 C) and the best performing constructs of all eight enzymes. All constructs contained either the promoter *P_H_p_FMD_* or *P_H_p_MOX_,* which are endogenous to *H. polymorpha.* We succeeded in transforming *H. polymorpha* and 96 well cultivation under comparable conditions as before with *P. pastoris*. Transformed episomal plasmids could be recovered and retransformed into *E. coli* and verified by sequencing, completing the standard episomal workflow of *P. pastoris*. Unfortunately, the protein production of the eight enzymes was not successful. No UPO/lipase/laccase activity was detected. These results point towards an incompatibility of the expression cassette, most likely caused by the utilized terminator (GAP terminator, *P. pastoris*) which might not be functional in *H. polymorpha* thereby impeding the protein production. The other expression unit parts (promoter, signal peptides, genes and protein-tag) should be functional per se.

## DISCUSSION

In the present study, we have developed an episomal *P. pastoris* expression system for combining strong methanol-inducible promoters derived from *P. pastoris* and *H. polymorpha* and a diverse signal peptide panel for target protein secretion. Modular Golden Gate cloning enables the rapid, effective construction of highly diverse promoter/signal peptide libraries with up to 187 unique combinations in a one-pot, one-step manner. This approach complements valuable existing systems, which provide useful Golden Gate parts and circuits for *Pichia pastoris.*^15, 17^ Our focus was on building a high-throughput system, relying on reproducible episomal expression, high-transformation efficiencies, and using biocatalysts of high relevance and challenging production as test set. We expect that this research further promotes the application of *P. pastoris* as host in directed evolution approaches, enabling the tailoring of diverse and post-translationally modified biocatalysts.

We solely relied on cultivation and screening in 96 well plates to demonstrate the system’s high throughput potential and screened approx. 3200 primary transformants of eight diverse enzymes. Well-performing episomal constructs were identified for all enzymes, and enzyme activities were increased further by subsequent carbon source screening. The screenings revealed significant differences in activity distribution even when using the identical promoter. This concludes that the construct cannot be assessed solely based on its promoter expression strength (weak, medium, strong) and regulation pattern. Secretion is substantially depending on the signal peptide/gene construct and its combination with a respective promoter. This insight points toward potential regulatory mechanisms during translation, protein folding, and glycosylation. A delicate balance, as high production rates of heterologous proteins can also overwhelm *P. pastoris’* cellular machinery and trigger protein degradation by unfolded protein response.^41^ Recent reports also suggest the occurrence of high amounts of retained non-secreted intracellular protein.^15^ Therefore, our system can provide a good starting point by supplying a diverse range of expression strength and signal peptides. Our strategy enables the rapid identification of constructs balancing high activity and high functionality. Based on the primary screen we could observe high (Mrl2: 70 %; *Tte*UPO: 66 %; *Mth*UPO: 56 %), medium (CalB: 44 %; *Mhi*UPO: 31% *Dca*UPO: 26 %; *Mfe*UPO 24 %) and low (AaeUPO*: 7 %) degrees of promiscuity towards occurring promoter/signal peptide combinations.

Our primary motivation was to develop a system to foster the discovery and subsequent engineering of known and novel UPOs, which are biocatalyst of outstanding interest but are also infamous for their challenging heterologous production.^2, 7–8^ We were able to optimize the episomal (3 of 3) and integrative production (2 of 3) of three known UPOs (MthUPO, TteUPO, *Aae-*UPO*). We successfully produced the new enzymes *Mhi*UPO and *Mfe*UPO for the first time. Also, the first production of *DcaUPO* in yeast was achieved, increasing the obtained yield of recombinant enzyme substantially. The overall volumetric yields of UPO production proved to be high, ranging from 5.7 to 22.4 mg/L. The designed two-layer system of compatible episomal and integrative plasmids enables the performance of the complete enzyme engineering workflow in *P. pastoris*. Following the identification of a suitable promoter-signal peptide combination and carbon source screening for obtaining maximal activity, the enzyme of choice can be rapidly evolved within the episomal system and interesting, final variants directly transferred to the integrative construct for subsequent high yield antibiotic-free protein production and characterization. Thereby we are bridging a gap in previous UPO engineering as well as directed evolution approaches in *P. pastoris* in general caused by the lack of suitable episomal plasmids for high efficiency transformation and reproducible expression. This limitation was previously overcome by a dual host approach, evolving the UPO in an episomal *S. cerevisiae* system and subsequently transferring the evolved variant for high yield protein production to *P. pastoris*. ^14^

Analysis of the distribution of promoter and signal peptide modules among the sequenced non-redundant top constructs (80 samples) provided several interesting insights (Supplemental Table 5). Signal peptide analysis revealed a broad distribution. The most frequently occurring signal peptides were “Sce-α Galactosidase” and “Gga-Lysozym” with 12.5 % as well as “Mro-UPO” and “Gma-UPO” with 11.3 %. Surprisingly especially signal peptides derived from UPOs (6 out of 17 in total; 33 % occurrence) proved to be highly valuable to target protein secretion in *P. pastoris*— also in case of the unrelated enzymes CalB and Mrl2. The α-factor leader “Sce-Prepro” is derived from *S.cerevisiae* and utilized as standard signal peptide for target protein secretion in yeast and included in nearly all commercial secretory *P. pastoris* plasmids. However, it was only identified in 3.8 % of our top hits. This further emphasizes the appeal of a one-pot, one-step signal peptide shuffling approach as it allows the probing of a diverse set of signal peptides for rapid secretion testing.

Besides the signal peptides, substantial new insights were gained into endogenous and orthologous promoters. Widely used promoters within integrative plasmids *(P_P_p_DAS1_; P_P_p_DAS2_; P_P_p_AOX1_)* exhibit strong repression in presence of glucose and glycerol and strong induction profiles upon methanol addition. *P_P_p_AOX1_* thereby is the most widespread promoter and is licensed in most commercial plasmid systems.^19, 42^ In our setup, these three promoters turned out to be the least occurring promoter among the top hits *(P_P_p_DAS1_; P_P_p_DAS2_:* 1.3 %; *P_P_p_AOX1_*: 2.5 %). We assume that these strongly regulated/induced promoters might be less suitable for the use in episomal systems based on their expression dynamics and strength. The most suitable promoters were *P_P_p_FLD1_* (23.8 %), *P_H_p_FMD_* (16.3 %) and *P_P_p_ALD4_* (15 %). In general, we observed that the derepression behavior is strongly correlating with the suitability and occurrence within the episomal system. Promoters which exhibited a pronounced activity under derepression conditions like *P_P_p_ALD4_, P_P_p_FDH1_, P_P_p_FLD1_, P_H_p_FMD_* and *P_P_p_CAT1_* (Figure 1 D), were present in 80 % of all top constructs. However, promoters with a strict methanol dependent induction and low derepression like *P_H_p_MOX_P_P_p_PMP20_, P_p_p_DAS1_, P_P_p_DAS2_* and *P_P_p_AOX1_* were the least often occurring promoters.

Astonishingly, for all enzymes the top episomal and integrative constructs contained orthologous promoter (6 x *P_H_p_FMD_;* 2 x *P_H_p_-MOX_*), the integrative strains often being two-to three-fold superior compared to the second or third best strains *P. pastoris* derived promoters (data not shown). Solely the third *H. polymorpha* derived promoter *P_H_p_DHAS_* did not stand out, however was found among within an average episomal distribution (6.3 %)—more often than *P_H_p_MOX_* (5 %).

These observations confirm the high potential of orthologous *H. polymorpha* promoters and further prove their high suitability on an episomal level. Utilizing these orthologous promoters and without performing any optimization on protein production of the diverse enzyme set, we were able to achieve mg/L yields of all enzymes in shake flask format. In most cases we could surpass previously reported yields of the respective enzymes. There is still potential of reaching higher volumetric yields after individual optimization of each protein production setup. *P_H_p_FMD_* proved to be an especially valuable promoter not just resulting in high induced yields of up to 30 mg/L but being also active under methanol-free conditions with 17 to 72 % of the maximal yield achieved under induced conditions. This feature renders methanol-free production a feasible procedure addressing safety and health concerns.

Pleasingly, the designed two-layer system yielded good results in both setups. It consists of an initial episomal assembly and screening (layer 1) and subsequent transfer of the entire expression unit to an integrative plasmid (layer 2). This approach led to the discovery of highly active episomal as well as integrative strains. Another scenario would have been the selection of expression units, which are functioning optimal for episomal production, but are less suitable as units for high-yield protein production when integrated into the genome. This hypothesis is also connected to the observed effect that the set of repressed/strong induced promoter *(P_P_p_DAS1_; P_P_p_DAS2_; P_P_p_AOX1_)* is clearly underrepresented within the top-performing episomal hits. However, if screening would have been conducted solely using an integrative system, the proportion of this promoters would have most likely increased due to their general high production rates when used as integrative construct.^42^

## CONCLUSION

The designed two-layer system of compatible episomal and integrative plasmids enables the performance of a complete enzyme engineering workflow in *P. pastoris*. Following the identification of a suitable promoter/signal peptide combination and carbon source activity screening, the enzyme of choice can be rapidly evolved within the episomal system—analogous to directed evolution approaches in *S. cerevisiae*. Interesting constructs or enzyme variants can be transferred directly to a matching integrative vector for high yield antibiotic-free protein production.

We are bridging a gap to previous UPO engineering and directed evolution approaches in *P. pastoris* in general. This gap was caused by the lack of suitable episomal plasmids. This limitation was previously overcome by a dual host approach, evolving the UPO in an episomal *S. cerevisiae* system and subsequently transferring the evolved mutant for high yield protein production to *P. pastoris.^14^* To highlight the versatility of the developed system, it was applied to optimize the episomal yield of two other enzyme classes: The lipase CalB and the laccase Mrl2. All relevant plasmids are available at Addgene.

## MATERIALS AND METHODS

### Bacterial and Yeast strains

For all cloning purposes and plasmid propagation *E. coli* DH10B cells (ThermoFisherScientific, Waltham, US) were utilized. All work regarding *Pichia pastoris* was performed utilizing the mut^+^ Strain X-33 (ThermoFisherScientific, Waltham, US). All work regarding *Hansenula polymorpha* was performed utilizing the wildtype strain Ha-1301 (DSMZ, Braunschweig, DE).

### Microtiter Plate cultivation expression of *P. pastoris* and H. *polymorpha*

For enzyme production in microtiter plate format specialized 96 half deep well plates were utilized. The model type CR1496c was purchased from EnzyScreen (Heemstede, NL) and plates were covered with fitting CR1396b Sandwich cover for cultivation. Plates and covers were flushed before every experiment thoroughly with 70 % ethanol and air dried under a sterile bench until usage. Each cavity was filled with 220 μL of buffered complex medium (BM) and inoculated with single, clearly separated yeast colonies using sterile toothpicks. Basic BM (20 g/L peptone; 10 g/L yeast extract; 100 mM potassium phosphate buffer pH 6.0; 1x YNB (3.4 g/L yeast nitrogen base without amino acids; 100 g/L ammonium sulfate); 400 μg/L biotin; 3.2 mM magnesium sulfate; 25 mg/L chloramphenicol; 50 mg/L hemoglobin; 150 mg/L Hygromycin B) was freshly prepared out of sterile stock solutions immediately before each experiment, mixed and added to the cavities.

Depending on the type of experiment different carbon source feeding strategies were employed. Therefore, defined amounts (0.3 %; 0.5 %; 1.0 % or 1.5 % final) of the primary carbon sources glucose and glycerol were added to the cultivation media derived from defined stock solutions. Pure methanol was added to a final concentration of 1.5 or 2 % (v/v). After inoculation of the wells the plates were covered, mounted on CR1800 cover clamps (EnzyScreen) and incubated in a Minitron shaking incubator (Infors, Bottmingen, SUI) for 72 h (30 °C; 230 rpm). After cultivation the cells were separated from the enzyme containing supernatant by centrifugation (3400 rpm; 50 min; 4 °C).

### Peroxygenase activity measurement via NBD assay

The use of 5-nitro-1,3-benzodioxole (NBD) as a suitable microtiter plate substrate for the measurement of peroxygenase catalysed conversion to the colorimetric product 4-Nitrocatechol has been described before.^11,34^ The described conditions have been adapted with slight modifications. In brief 20 μL or 40 μL of peroxygenase containing supernatant (from 96 well plate setup) are transferred to a transparent polypropylene 96 well screening plate (Greiner Bio-One, Kremsmünster, AT) and 160 or 180 μL of screening solution (final: 100 mM potassium phosphate pH 6.0; 1 mM NBD; 1 mM hydrogen peroxide; 12 % (v/v) acetonitrile) added. Absorption values (λ: 425 nm) of each well were immediately measured after addition and brief shaking (3 sec) in a kinetic mode (measurement interval: 30 s) over a duration of 5 to 20 minutes utilizing the 96 well microtiter plate reader Spark 10M (TECAN, Grödig, AT). Slope values of absorption increase corresponding to 4-nitrocatechol formation were obtained, paying special attention to the linearity of the observed slope to obtain reliable relative NBD conversion values for comparison of the respective wells.

In the case of the initial screening of novel peroxygenases the screening conditions were slightly modified. 40 μL of supernatant were used and NBD as well as H2O2 concentrations reduced to 300 μM to prevent solubility issues. 4-nitrocatechol formation was monitored as previously described for 45 minutes.

### Peroxygenase activity measurement via DMP assay

Rescreening setups of novel peroxygenases were performed utilizing 2,6-Dimethoxyphenol (DMP) as microtiter plate substrate. The use of DMP as suitable substrate for the measurement of peroxygenase catalyzed conversion to the colorimetric product ceruglinone has been described before.^12^ The described conditions have been adapted with slight modifications. In brief, 40 μL of peroxygenase containing supernatant (from 96 well plate setup) are transferred to a transparent polypropylene 96 well screening plate (Greiner Bio-One, Kremsmünster, AT) and 160 μL of screening solution (final: 100 mM potassium phosphate pH 6.0; 3 mM 2,6-Dimethoxyphenol; 1 mM hydrogen peroxide) added. Absorption values (λ: 469 nm) of each well were immediately measured after addition in a kinetic mode (measurement interval: 30 s) over a duration of 15 minutes utilizing the 96 well microtiter plate reader Spark 10M (TECAN, Grödig, AT). Slope values of absorption increase corresponding to ceruglinone formation were obtained, paying special attention to the linearity of the observed slope to obtain reliable relative DMP conversion values for comparison of the respective wells.

### CalB lipase activity measurement via 4-Nitrophenyl laurate conversion

40 μL of lipase containing supernatant were transferred to a transparent polypropylene 96 well screening plate (Greiner Bio-One, Kremsmünster, AT) and 160 μL screening solution (final: 100 mM Tris-HCl pH 8.0; 1 mM 4-Nitrophenyl laurate; 1 % (v/v) Triton X-100) added. Absorption values (λ: 405 nm) of each well were immediately measured after addition and brief shaking (3 sec) in a kinetic mode (measurement interval: 30 s) over a duration of 15 minutes utilizing the 96 well microtiter plate reader Spark 10M (TECAN, Grödig, AT). Slope values of absorption increase corresponding to 4-Nitro-phenolat release were obtained, paying special attention to the linearity of the observed slope to obtain reliable relative 4-Nitro-phenyl laurate conversion values for comparison of the respective wells.

### Mr2 laccase activity measurement based on ABTS/DMP conversion

10 μL of laccase containing supernatant were transferred to a transparent polypropylene 96 well screening plate (Greiner Bio-One, Kremsmünster, AT) and 190 μL screening solution (final: 100 mM sodium citrate pH 4.0; 100 μM ABTS). Absorption values (λ: 418 nm) of each well were immediately measured after addition and brief shaking (3 sec) in a kinetic mode (measurement interval: 30 s) over a duration of 10 minutes utilizing the 96 well microtiter plate reader Spark 10M (TECAN, Grödig, AT). Slope values of absorption increase corresponding to the formation of the radical ABTS cation (ABTS•+) were obtained, paying special attention to the linearity of the observed slope to obtain reliable relative ABTS conversion values for comparison of the respective wells.

Due to the rapid conversion of ABTS by Mrl2 later screenings were performed using DMP as substrate to obtain reliable slope values. Therefore 10 μL of supernatant and 190 μL of screening solution (final: 100 mM potassium phosphate pH 6.0; 100 μM DMP) was utilized and measurement performed as described above.

## ASSOCIATED CONTENT

All crucial plasmids (17 x signal peptides; 4x episomal plasmids; 1x integrative plasmid; 11 × MUT promoters and 7× C-terminal protein tags etc.) have been deposited as comprehensive, modular Kit with the non-profit plasmid repository Addgene (Yeast Secrete and Detect).

Detailed information regarding experimental procedures as well as additional data can be found within the supplemental information. This material is available free of charge via the Internet at http://pubs.acs.org.

## AUTHOR INFORMATION

### Author Contributions

P.P. envisioned and constructed the modular secretion system.

P.P. planned the research and conducted the experiments. P.P. and M.J.W. wrote the manuscript.

## ACKNOWLEDGMENT

M.J.W thanks the Bundesministerium für Bildung und Forschung („Biotechnologie 2020+ Strukturvorhaben: Leibniz Research Cluster”, 031A360B and 031A360E) for generous funding. P.P. thanks the Landesgraduiertenförderung Sachsen-Anhalt for a PhD scholarship. We would like to thank Cătălin Voiniciuc (Leibniz Institute of Plant Biochemistry; IPB Halle) for providing Pichia plasmid parts and the X33-strain. We are especially grateful towards Dr. Falko Matthes (IPK Gatersleben) for his rapid and generous gift of genomic DNA of *Hansenula polymorpha.* Michael Niemeyer (IPB Halle) and Jann Simon Groen (MLU Halle) are acknowledged for advice and support regarding protein purification.

## ABBREVIATIONS

MUT: methanol utilization pathway
ARS: autonomously replicating sequence
AOX1: alcohol oxidase 1
FDH1: Formate Dehydrogenase 1
FLD1: Formaldehyde dehydrogenase 1
PMP20: Peroxisomal glutathione oxidase 20
DAS1: Dihydroxyacetone synthase 1
DAS2: Dihydroxyacetone synthase 2
CAT1: Catalase 1
ALD4: Mitochondrial aldehyde dehydrogenase 4
DHAS: Dihydroxyacetone synthase
FMD: Formate dehydrogenase
MOX: methanol oxidase
Gly: glycerol
NBD: 5-nitro-1,3-benzodioxole
DMP: 2,6-Dimethoxyphenol
IEX: ion exchange
HIC: hydrophobic interaction chromatography
MW: molecular weight
ABTS: 2,2’-azino-bis(3-ethylbenzothiazoline-6-sulfonic acid
GAP: glyceraldehyde-3-phosphate dehydrogenase

## Table of contents graphic

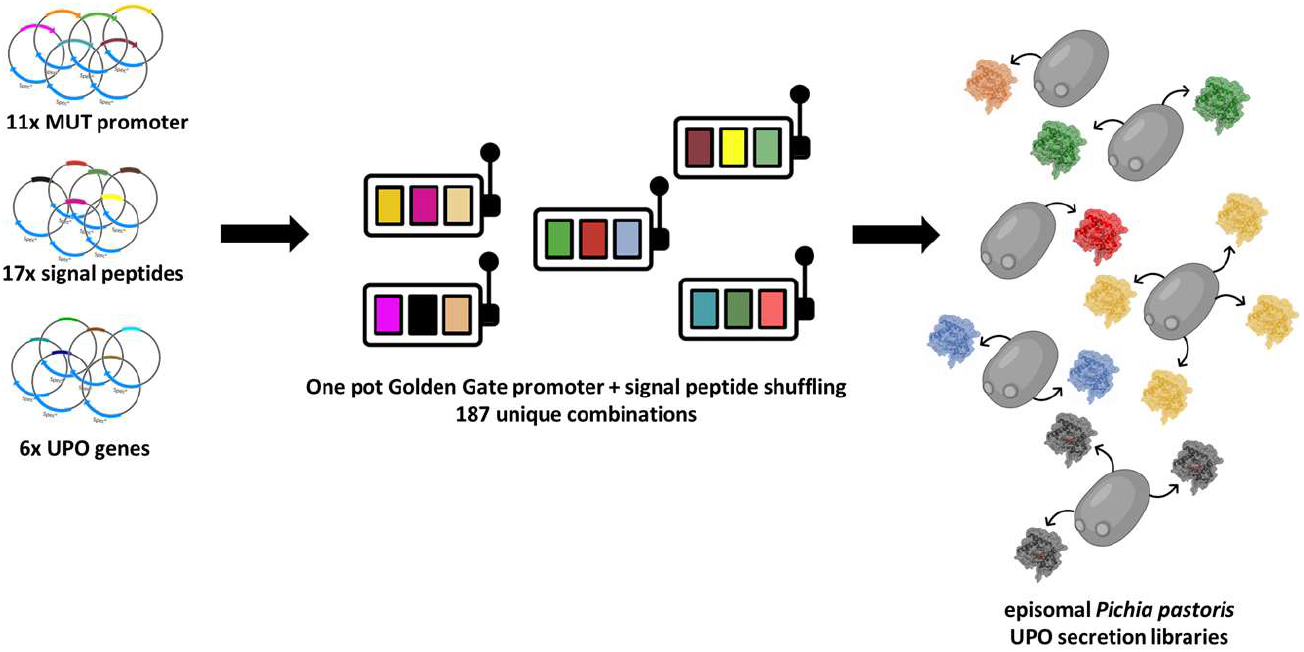

