## Supplemental information for "Improving the heterologous production of fungal peroxygenases through an episomal *Pichia pastoris* promoter and signal peptide shuffling system"

### Material and Methods

**Chemicals.** Solvents were used as received without further purification. Acetone was utilised as GC ultra-grade (≥ 99,9 %) from Carl Roth (Karlsruhe, DE). Acetonitrile was purchased from Merck (Darmstadt, DE) in gradient grade for LC (≥ 99,9 %). Methanol (LC-MS grade ≥ 99,9 %) was used from Honeywell Chemicals (Seelze, DE). 5-Nitro-1,3-benzodioxole (NBD; 98 % purity) and 2,2′-Azino-bis(3-ethylbenzothiazoline-6-sulfonic acid) diammonium salt (ABTS) were retrieved from Sigma-Aldrich (Hamburg, DE). 4-Nitrophenyl laurate was purchased from TCI Chemicals (Tokyo, JP).

**Enzymes and cultivation media.** For cultivation of *E. coli* cells terrific broth (TB) media from Carl Roth (Karlsruhe, DE) was used. Peptone and Yeast extract were purchased from Formedium (Hunstanton, GB). Yeast nitrogen base (without amino acids) and Hygromycin B solution (50 mg/ml) were purchased from Carl Roth (Karlsruhe, DE). PNGase F and BsaI were purchased from New England Biolabs (Ipswich, US). BbsI and FastDigest AscI were purchased from ThermoFisherScientific (Waltham, US) and T4 DNA Ligase from Promega (Madison, US).

### General Procedures

**Expression plasmid construction (pPAP004) for *Pichia pastoris.*** The design of the episomal backbone plasmid pPAP004 is based on a previously constructed Pichia system consisting of two episomal expression plasmids (pPAP001 and pPAP002) and an integrative plasmid (pPAP03) for subsequent stable genomic integration.[1] The level 1 Golden Gate based shuttle expression plasmid pPAP004 was therefore PCR amplified (Phusion Polymerase) from pPAP001 excluding the pGAP promoter and assembled by Golden Gate cloning using BsmBI. It can be propagated in *E. coli* (Ampicillin resistance) as well as *P. pastoris* (Hygromycin B resistance). To enable episomal plasmid propagation in *P. pastoris,* the plasmid is equipped with a previously described functional ARS sequence, which was originally PCR amplified from *Kluveromyces lactis* genomic DNA.[2-3]
As placeholder for a target gene sequence as well as color selection marker, a lacZ cassette (approx. 600 bp) is integrated. Upon digestion with BsaI the lacZ cassette is released and a fitting open reading can be integrated in frame (e.g. promoter-Signal Peptide-Gene-C-terminal Tag) into the plasmid, thereby reconstituting a fully functional expression plasmid. For the stable integration of transcription units into the genome of *P. pastoris,* a third universal integrative plasmid (pPAP003) was utilized. This shuttle plasmid was previously constructed, which can be propagated in *E. coli* (Kanamycin resistance) as well as *P. pastoris* (Hygromycin B resistance). As placeholder for a target transcription unit a lacZ cassette is integrated, enabling blue/white selection of transformants based on the conversion of X-Gal. Upon digestion with BbsI the lacZ cassette is released and a fitting transcription unit (Promoter- ORF- Terminator) can be integrated (derived from respective pPAP004 episomal plasmids as donors) into the plasmid, thereby reconstituting a fully functional integration plasmid which can then be used for transformation.

**Construction of Level 0 parts.** Individual assembly parts (genes and promoters) were cloned as individual parts into the respective pAGM9121 Level 0 universal acceptor plasmid. All new gene parts (CalB, Mr2 Laccase, *Mfe*UPO, *Mhi*UPO and *Dca*UPO) were purchased as double stranded DNA parts from Twist Bioscience (San Francisco, US), codon optimized for *Pichia pastoris*, bearing terminal BbsI recognition sites and pre-designed overhangs for correct assembly into pAGM9121.
In the case of the construction of the promoter library different strategies were employed. All promoter parts were cloned as individual parts into the respective pAGM9121 Level 0 and contain the same Kozak Sequence ($^{-10}$ATCATACAAA$^{0}$**ATG**). Several promoter parts ($P_{PpAOX1}$, $P_{PpFDH1}$, $P_{Pp\mathrm{DAS1}}$, $P_{Pp\mathrm{DAS2}}$, $P_{Pp\mathrm{ALD4}}$) were PCR amplified and derived from a previously introduced Golden Gate based *P. pastoris* assembly system, coined GoldenPics.[4] These parts are particularly suitable since they have been already domesticated regarding internal BsaI and BbsI recognition sequences. $P_{Pp\mathrm{FLD1}}$ and $P_{Pp\mathrm{PMP20}}$ were PCR amplified from genomic *Pichia pastoris* DNA. The $P_{Hp\mathrm{MOX}}$ element was amplified from genomic *Hansenula polymorpha* DNA in a comparable manner. Since the genomic sequences of $P_{Hp\mathrm{FMD}}$ and $P_{Hp\mathrm{DHAS}}$ contain internal recognition sites, both parts were purchased as double stranded DNA parts from Twist Bioscience (San Francisco, US) as described above and internal BsaI/BbsI recognition sites excluded by point mutagenesis. All Level 0 parts were constructed by Golden Gate digestion ligation reactions (+BbsI and T4-Ligase) using pAGM9121 as acceptor plasmid. All parts were subsequently confirmed by Sanger sequencing.

**Golden Gate Cloning of Expression Plasmids.** The expression plasmid pPAP004 was used as respective acceptor plasmid for the assembly of the individual tetrapartite expression units (**5′** promoter-Signal Peptide-Gene-C-terminal Tag **3′**). The individual parts were thereby derived as parts from standard level 0 plasmids (pAGM9121 backbone), which can be released from the pAGM9121 backbone upon BsaI restriction digest.
Golden Gate reactions were performed in a total volume of 15 µL. The final reaction volume contained 1-fold concentrated T4 ligase buffer. Prepared reaction mixtures containing ligase buffer, the acceptor plasmid (20 fmol) and the corresponding inserts as level 0 modules (Promoter, Signal Peptide, Gene, C-terminal Tag) were added to 20 fmol each and the overall volume adjusted to 13.5 µL with dd$H_2O$. In the case of shuffling approaches 17 different pAGM9121- Signal Peptide combinations were added in equimolar ratios (1.2 fmol each) and 11 different pAGM9121- promoter combinations in an equimolar ratio (1.9 fmol each) as well. In a final step, the corresponding enzymes were quickly added. First, a volume of 0.5 µL of the restriction enzyme BsaI (10 units/µL) was added and then 1 µL (1–3 units/µL) of T4 ligase. Golden Gate reactions where performed using a temperature cycling program (50 to 99x passes) between 37 °C (2 min) and 16 °C (5 min) and concluded by an additional enzyme inactivation step (80 °C; 20 min).
The whole Golden Gate reaction volume was used to transform 100 µL of chemically competent *E. coli* DH10B cells. After heat shock transformation (42°C; 90 sec) and recovery the mixture (approx. 320 µL) was split into two fractions, 50 µL were plated on selective LB

Agar plates (50 µg × $mL^{-1}$ X-Gal; 100 µg × $mL^{-1}$ Ampicillin; 150 µM IPTG) and the remaining volume used to directly inoculate 4 mL TB Medium (100 µg × $mL^{-1}$ Ampicillin) to preserve the genetic diversity of the shuffling library. The following day the success of the Golden Gate reaction was evaluated based on the performed blue/white screening, discriminating the empty plasmid (lacZ; blue) from recombined, white colonies. In general, the described protocol for ORF assembly and promoter + signal peptide shuffling as special case led to several hundred recombinant colonies with a high proportion (> 90 %) of recombined, white colonies.
In the case of single defined, "unshuffled" constructs single colonies were checked for correct insert sizes by the means of colony PCR (matching pPAP004 sequencing primer). Positively identified clones were inoculated into 4 mL of TB-Medium (100 µg × $mL^{-1}$ Ampicillin) and corresponding plasmid DNA prepared (NucleoSpin Plasmid Kit (Macherey-Nagel, Düren, DE)). In the case of shuffled signal peptide constructs, plasmid DNA was prepared as a library by direct inoculation of the transformation mixture into liquid culture and subsequent DNA isolation (see above).

**Plasmid transformation into *P. pastoris*.** Respective single plasmids or plasmid mixtures (pPAP004 backbone) were used to transform electro competent *P. pastoris* cells (X-33 strain) by the means of electroporation. Electro competent X-33 cells were prepared according to a condensed protocol for *P. pastoris*.[5] Cells were stored in BEDS solution (10 mM bicine-NaOH pH 8.3, 3 % (v/v) ethylene glycol, 5 % (v/v) DMSO and 1 M sorbitol) as 40 µL aliquots (-80 °C) till further use.
For the transformation of episomal plasmids 60 ng of the circular plasmid (predilution in dd$H_2O$ to a 60 ng/µL stock solution) were added to one aliquot of on ice thawed competent X-33 cells. The cell-plasmid mix was then transferred to an electroporation cuvette (2 mm gap) and the cuvette cooled for 10 minutes on ice prior to the transformation. Electroporation was performed using a Micropulser Device (Bio-Rad, Hercules, US) and using manual implemented, standardized settings (1.5 kV, 1 pulse) for all transformation setups, leading to a general pulse interval of 5.4 to 5.8 ms. Immediately after electroporation cells were recovered in 1 mL of ice cold YPD-Sorbitol solution (10 g/L peptone, 5 g/L yeast extract, 500 mM sorbitol), transferred to a new tube and incubated for one hour under rigid shaking (30 °C, 900 rpm) in a Thermomix device (Eppendorf, Hamburg, DE). After incubation cells were precipitated by centrifugation (5700 rpm, 5 min). The supernatant was discarded, and the cells resuspended in 200 µL of fresh YPD medium. 100 µL of the suspension were then plated on selective YPD Agar plates supplemented with 150 µg/mL Hygromycin B. Plates were incubated at 30 °C for at least 48 h till clearly visible colonies appeared. In general, the described setup led to the occurrence of several hundred colonies per plate.
For the transformation of integrative plasmids (pPAP003 backbone) the setup was slightly modified since in this case linearized plasmids are used for transformation. Therefore, previously prepared circular plasmid DNA was digested with AscI (Isoschizomer: SgsI). 2.5 µg of the respective plasmid DNA was mixed with 3 µL of 10x fold FastDigest Buffer, the volume adjusted to 29.5 µL using dd$H_2O$ and in a last step 0.5 µL of FastDigest SgsI added. Digestion was performed overnight (16 h, 37 °C) and terminated by an enzyme inactivation step (20 min, 80 °C). Linearized plasmid DNA was then subsequently prepared according to the manufacturer instruction using a Nucleospin® Gel and PCR clean up Kit (Macherey-Nagel, Düren, DE). The transformation of *P. pastoris* was performed in a congruent manner as described before, except for using 100 ng linearized plasmid for transformation, since the overall transformation efficiency is substantially reduced in comparison to the transformation of circular plasmid.

**Plasmid transformation into *H. polymorpha*.** Respective single plasmids (pPAP004 backbone) were used to transform electro-competent *H. polymorpha* cells (X-33 strain) by the means of electroporation. Cells were prepared and transformed according to a previously published protocol.[6] After transformation the whole volume was plated on selective YPD Agar plates supplemented with 150 µg/mL Hygromycin B. Plates were incubated at 37 °C for at least 48 h till clearly visible colonies appeared.

**Microtiter Plate cultivation expression of *P. pastoris* and *H. polymorpha*.** For enzyme production in microtiter plate format specialized 96 half deep well plates were utilized. The model type CR1496c was purchased from EnzyScreen (Heemstede, NL) and plates were covered with fitting CR1396b Sandwich cover for cultivation. Plates and covers were flushed before every experiment thoroughly with 70 % ethanol and air dried under a sterile bench until usage. Each cavity was filled with 220 µL of buffered complex medium (BM) and inoculated with single, clearly separated yeast colonies using sterile toothpicks. Basic BM (20 g/L peptone; 10 g/L yeast extract; 100 mM potassium phosphate buffer pH 6.0; 1x YNB (3.4 g/L yeast nitrogen base without amino acids; 100 g/L ammonium sulfate); 400 µg/L biotin; 3.2 mM magnesium sulfate; 25 mg/L chloramphenicol; 50 mg/L hemoglobin; 150 mg/L Hygromycin B) was freshly prepared out of sterile stock solutions immediately before each experiment, mixed and added to the cavities.
Depending on the type of experiment different carbon source feeding strategies were employed. Therefore, defined amounts (0.3 %; 0.5 %; 1.0 % or 1.5 % final) of the primary carbon sources glucose and glycerol were added to the cultivation media derived from defined stock solutions. Pure methanol was added to a final concentration of 1.5 or 2 % (v/v).
Regarding the production of Mr2 laccase and CalB some alterations regarding the cultivation media were made. CalB expression plates were prepared without the supplementation of magnesium sulphate and hemoglobin. In the case of Mr2 Laccase expression plates 3.2 mM magnesium sulfate were replaced by 1 mM copper sulfate and no hemoglobin added. Volume differences were adjusted with dd$H_2O$ and the overall procedure not changed. In the case of rescreen setups of the novel peroxygenases (*Mhi*UPO, *Mfe*UPO and *Dca*UPO) medium was prepared without the supplementation of hemoglobin to avoid interference (background activity) within the utilized screening assay.
After inoculation of the wells the plates were covered, mounted on CR1800 cover clamps (EnzyScreen) and incubated in a Minitron shaking incubator (Infors, Bottmingen, SUI) for 72 h (30 °C; 230 rpm). After cultivation the cells were separated from the enzyme containing supernatant by centrifugation (3400 rpm; 50 min; 4 °C).

**Shake flask cultivation *P. pastoris*.** For large scale protein production using shake flasks genomically integrated single constructs (pPAP 003 backbone; integration into chromosomal 3′ region of *P. pastoris* AOX1 gene) were chosen. These constructs were previously identified by screening at least 7 different colonies per individual construct (promoter - signal peptide - gene combination) within a MTP screening setup and choosing a respective production strain based on a high as possible, clearly distinguishable NBD, ABTS or 4-nitrophenyl laurate conversion in comparison to the background control (pPAP003 empty plasmid).
Precultures were prepared in 50 mL YPD medium (+ 25 mg/L chloramphenicol) and cultivated for 48 h (30 °C; 160 rpm; 80 % humidity), typically resulting in a final $OD_{600nm}$ of approx. 17 to 19. The main expression culture was inoculated with a starting optical density of 0.3. For large scale production BM based expression media (20 g/L peptone; 10 g/L yeast extract; 100 mM potassium phosphate buffer pH 6.0;

1x YNB (3.4 g/L yeast nitrogen base without amino acids; 10 g/L ammonium sulfate); 400 µg/L biotin; 3.2 mM magnesium sulfate (peroxygenases) or 1 mM copper sulfate (Mr2 Laccase); 25 mg/L chloramphenicol) was utilized. Cultivation was performed in 2,5 L Ultra yield flasks (Thomson Instrument, Oceanside, US) in a final culture volume of 500 mL per flask after sealing the flask with breathable Aeraseal tape (Sigma Aldrich, Hamburg, DE) allowing for gas exchange. The main cultures were incubated for further 72 h (25 °C; 110 rpm; 80 % humidity).
In the case of derepressed production 1 % (w/v) Glycerol were added as carbon source for Pichia growth. In the case of the methanol induced production a two-phase feeding was applied, firstly inoculating the cells into BM medium (see above) supplemented with 0.4 % (w/v) glycerol as carbon source. 24 h and 48 h after inoculation 1.0 % (v/v) of methanol were added as inducer of the respective MUT promoter. After cultivation the cells were separated from the enzyme containing supernatant by centrifugation (4300 rpm; 35 min; 4 °C).

**Supernatant ultrafiltration and protein purification.** The previously prepared supernatant was concentrated approx. 20-fold by the means of ultrafiltration. Therefore, a Sartocon Slice 200 membrane holder (Sartorius, Göttingen, DE) was equipped with a Sartocon Slice 200 ECO Hydrosart Membrane (10 kDa nominal cut-off; Sartorius) within a self-made flow setup. The flow system for ultrafiltration was operated by an EasyLoad peristaltic pump (VWR International, Darmstadt, DE).
In a first step the cleared supernatant (1 L) was concentrated approx. 10-fold to a volume of 100 mL and 900 mL of purification binding buffer (100 mM Tris-HCl pH 8.0, 150 mM NaCl) were added as a buffer exchange step. This sample was then concentrated approx. 20-fold to achieve a final volume of 50 mL.
Protein purification was implemented utilizing the C-terminal attached double Strep II Tag (WSHPQFEK), coined TwinStrep® (Iba Lifesciences, Göttingen, DE). As column material Strep-Tactin®XT Superflow® columns (1 mL or 5 mL; Iba Lifesciences) were chosen and the flow system operated by an EasyLoad peristaltic pump (VWR). In a first step the column was equilibrated with 5 column volumes (CVs) binding buffer. The concentrated sample (50 mL) was filter sterilized (0.2 µm syringe filter) and applied to the column with an approximate flow rate of 1 mL/min. After application the column was washed with 6 CVs binding buffer. Elution was performed based on binding competition with biotin, therefore approx. 2 CV of elution buffer (100 mM Tris-HCl pH 8.0, 150 mM NaCl; 50 mM biotin) were applied to the column. The pooled elution fraction was then dialysed overnight (4 °C) against 5 L of storage buffer (100 mM potassium phosphate pH 7.0) using ZelluTrans dialysis tubing (6-8 kDa nominal cut-off; Carl Roth) and the recovered, dialyzed sample stored at 4 °C till further use.

**Purification of *Dca*UPO by hydrophobic interaction chromatography (HIC).** The enzyme *Dca*UPO was purified by the means of hydrophobic interaction chromatography (HIC). Therefore, the cultivation supernatant was first concentrated as described above to a volume of approx. 100 mL and approx. 950 mL of equilibration buffer (50 mM potassium phosphate; pH 7.0) added. The sample was concentrated once again to a final volume of 50 mL and filter sterilized (0.2 µm syringe filter). Afterwards a concentrated ammonium sulfate solution (3.5 M stock solution) was slowly added using a syringe pump for dosing till reaching a final concentration of 1 M within the sample after 40 min under stirring. Precipitated proteins were separated by centrifugation (12000 rpm; 15 min; 20 °C) from the soluble fraction. A 1 mL HiTrap™ Octyl FF column was used as material (GE Healthcare, Uppsala, SWE) and equilibrated with binding buffer (33 mM potassium phosphate pH 7.0; 1 M $NH_4SO_4$). The sample was applied to the column with an approximate flow rate of 1 mL/min and the column washed afterwards with approx. 20 CV of binding buffer. Primary elution was achieved with a sharp step elution by switching to low salt elution buffer (33 mM potassium phosphate pH 7.0) and approx. 4 CV of elution fraction retrieved. A tightly bound fraction of *Dca*UPO remained on the column and was then eluted as highly pure enzyme using pure ethylene glycol as eluent. The ethylene glycol elution fraction was then dialyzed overnight (4 °C) against 5 L of storage buffer (100 mM potassium phosphate pH 7.0) using ZelluTrans dialysis tubing (6-8 kDa nominal cut-off; Carl Roth) and the recovered, dialyzed sample stored at 4 °C till further use.

**Plasmid preparation of episomal plasmids from yeast.** Yeast plasmids of identified clones were recovered by the means of digestive Zymolase cell treatment and alkaline cell lysis. Therefore streak-outs of previously identified suitable promoter-signal peptide combinations on YPD-Agar plates were used. A small amount of cells was collected with a sterile toothpick and resuspended in 1 mL of washing buffer (10 mM EDTA NaOH; pH 8.0) by light vortexing. Cells were subsequently pelleted (5000 × g; 10 min) and the supernatant discarded. Afterwards cells were resuspended by light vortexing in 600 µL of Sorbitol Buffer (1.2 M sorbitol, 10 mM $CaCl_2$, 100 mM Tris-HCl pH 7.5, 35 mM β-mercaptoethanol) and 100 units of Zymolase (Sigma Aldrich, Hamburg, DE) added followed by an incubation step for 120 min (30 °C; 650 rpm) to facilitate yeast cell wall digestion. After incubation cells were pelleted by centrifugation (2000 × g; 10 min) the supernatant discarded, and the plasmid preparation started with an alkaline lysis step following the manufacturer's instructions (NucleoSpin Plasmid Kit, Macherey Nagel). In a final step the yeast derived episomal plasmids were eluted in 25 µL elution buffer (5 mM Tris-HCl pH 8.5) and the whole eluate used to transform one aliquot of *E. coli* DH10B (transformation as described above), plating the whole transformation mix on a selective LB-Agar plate (50 µg × $mL^{-1}$ X-Gal; 100 µg × $mL^{-1}$ Ampicillin; 150 µM IPTG).
On the following day single colonies were picked, inoculated into 4 mL of TB medium (100 µg × $mL^{-1}$ Ampicillin), plasmid prepared (NucleoSpin Plasmid Kit) and sent for Sanger Sequencing to elucidate the respective sequence of the open reading frame (Eurofins Genomics).

**Protein concentration determination and purification yield.** Protein concentrations of the respective protein samples were determined after dialysis of the elution fractions (storage buffer: 100 mM potassium phosphate pH 7.0). In this regard the colorimetric BCA assay was utilized, employing a Pierce™ BCA Protein Assay Kit (ThermoFisherScientific, Waltham, US) following the instructions of the manufacturer. Samples were measured in biological triplicates (25 µL of a previously diluted sample) and concentrations calculated based on a previously performed calibration curve using BSA (0 - 1000 µg/mL) as reference protein.
To determine the overall yield of enzyme production per litre of culture volume the determined concentration in the elution fraction was extrapolated to the overall NBD/ABTS/4-Nitrophenyl laurate activity of the sample after ultrafiltration (column load). This calculation is based on the fact that all substrates are suitable for specific activity measurement, since negligible background signals of empty plasmid control (pPAP003 and pPAP004) regarding the substrate's conversions were observed.
Therefore, samples of every purification step (load, flow-through, wash and elution fraction) were collected and NBD/ABTS/4-Nitrophenyl laurate conversion rates of the respective fractions measured immediately after purification by testing of suitably diluted samples. In the case of non-complete binding of the enzyme fraction to the affinity column (remaining enzymatic activity in flow-through fractions) this remaining non-bound enzyme amount was taken into consideration for calculation for the overall volumetric production yield per litre. Therefore,

the via BCA assay determined protein concentration of the elution fraction was extrapolated to the activity of the respective non-bond fraction, assuming a comparable specific enzyme activity for NBD/ABTS/4-Nitrophenyl laurate conversion and considering the volumes of the respective fractions, leading to an approximate enzyme titre per litre of shake flask culture.

**Resting-state absorption and heme CO complex measurements.** The pooled and dialyzed elution fractions (100 mM potassium phosphate pH 7.0) were used to record absorption spectra of the respective enzymes in their native, resting state (ferric iron; $Fe^{3+}$). For all measurements a QS High precision Quartz Cell cuvette (Hellma Analytics, Müllheim, DE) with a path length of 10 mm was used. Spectra were recorded on a Biospectrometer Basic device (Eppendorf, Hamburg, DE) in the spectral range from 250 to 600 nm (interval: 1 nm) and subtracting the utilized storage buffer (100 mM potassium phosphate pH 7.0) as previous blank measurement.
Heme carbon dioxide spectra (CO assay) were recorded after reducing the heme iron to its ferrous form ($Fe^{2+}$). Therefore, a spatula tip of sodium dithionite as reducing agent was added to 1 mL of a respective enzyme fraction (see above) and mixed thoroughly till complete dissolution. This sample was split into two fractions and one fraction immediately flushed with a constant carbon monoxide flow for 2 min (approx. 1 bubble/sec) to obtain the thiolate-heme carbon monoxide complex. The sample was immediately transferred to a cuvette and absorption measured as described above, subtracting the reduced non-treated sample as blank measurement.
In case of *Mfe*UPO and *Mhi*UPO CO-Assay was also used to quantify volumetric yields of UPO production in comparison with *Mth*UPO. Therefore, a spatula tip of sodium dithionite as reducing agent was added to 1 mL of the enriched (approx. 10x fold) respective enzyme fraction (see above) after ultrafiltration (in 100 mM potassium phosphate pH 6.0) and mixed thoroughly till complete dissolution. This sample was split into two fractions and one fraction immediately flushed with a constant carbon monoxide flow for 2 min (approx. 1 bubble/sec) to obtain the thiolate-heme carbon dioxide complex. The sample was immediately transferred to a cuvette and absorption measured as described above, subtracting the reduced non-treated sample as blank measurement. pPAP003 was measured as a negative control, exhibiting no detectable maximum around 450 nm. Due to the high sequence homology to *Mth*UPO (> 90 %) the volumetric yield of *Mth*UPO (22.4 mg/L) was set as 100 % and both enzyme yields (*Mfe*UPO and *Mhi*UPO) calculated based on their maximal peak intensity when compared to the peak intensity of a MthUPO sample which was prepared under identical conditions.

**SDS Gel analysis and PNGaseF treatment.** Obtained elution fractions of the respective enzymes were analyzed for the apparent molecular weight and overall purity after the performed one step TwinStrep purification by the means of SDS PAGE. Therefore, samples of the column load (after ultrafiltration; see above), elution fractions after dialysis and deglycosylated elution fraction samples were analyzed on self-casted SDS PAGE (10 or 12 % of acrylamide) utilizing a Bio-Rad (Hercules, US) Mini-Protean® Gel electrophoresis System. For the purpose of molecular weight determination, a defined PageRuler Prestained Protein Ladder (ThermoFisherScientific, Waltham, US) was included, covering a MW range between 10 and 180 kDa. Proteins were visualized using a colloidal Coomassie G-250 staining solution.
To obtain N-type deglycosylated protein samples elution fractions were enzymatically treated with Peptide-N-Glycosidase F (PNGaseF) from *Flavobacterium meningosepticum*, which is capable of cleaving Asparagine linked high mannose type glycan structures as typically occurring in P. pastoris derived glycosylation patterns. Therefore, 45 µL of a respective elution fraction were mixed with 5 µL of denaturing Buffer (final 0.5 % SDS; 40 mM DTT) and denatured for 10 minutes (100 °C). After a cooling step to room temperature 6 µL of NP-40 solution (final: 1 %) and 6 µL of GlycoBuffer2 (500 mM sodium phosphate; pH 7.5) were added and the solution thoroughly mixed. Finally, 1 µL of PNGaseF (New England Biolabs, Ipswich, US) was added and the sample incubated under light shaking (37 °C) for 3 hours. After incubation the sample was prepared for further analysis by adding 5x fold SDS sample buffer and subsequent SDS PAGE analysis executed as described before.

**Supplemental Figure 1 General design principle of Golden Gate Assembly**

Schematic overview of designed Golden Gate circuit. (1) Individual modules (promoter, signal peptide, UPO gene,C terminal Tag) are cloned as Level 0 modules into pAGM9121 and flanked by BsaI recognition sites. Upon BsaI digest parts are released bearing the indicated 4 bp overhangs for subsequent reassembly. (2) The tetrapartite assembly can be correctly assembled into pPAP004 (BsaI digest). (3) Whole transcription units (promoter-ORF-terminator) can be swapped into a genomic integration plasmid (BbsI digest).

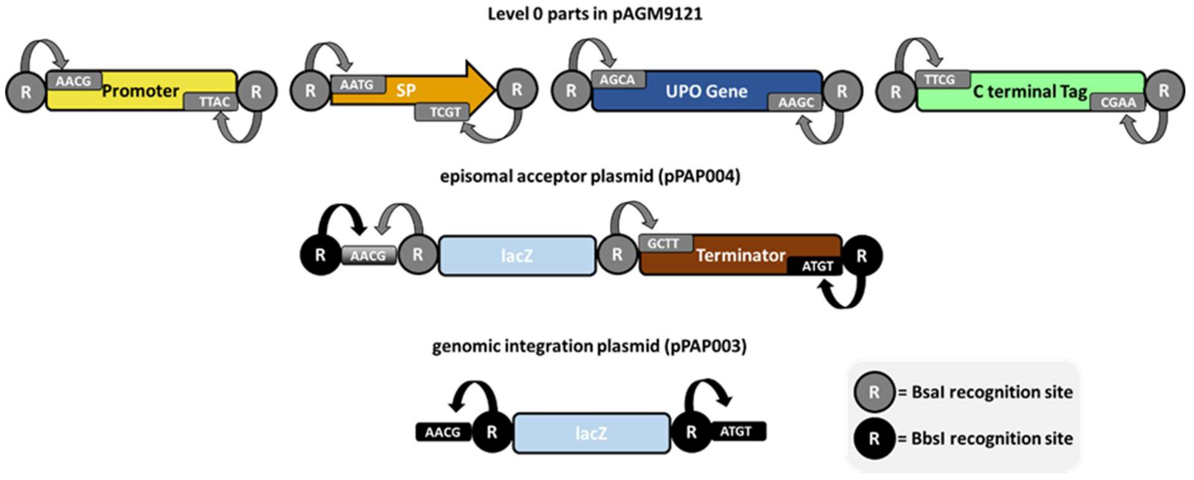


**Supplemental Figure 2 Absolute production levels *Tte*UPO utilizing different promoters and carbon source feeds**

Respective constructs ($P_X$- *Sce*−Invertase 2 SP-*Tte*UPO) were cultivated in 96 well plates for 72 h and samples (6 biological replicated) measured for UPO activity. The highest mean of NBD conversion ($P_{CAT1}$; 0.5 % Glycerol and 1.5 % methanol) was set as 100 % and all values normalized accordingly.

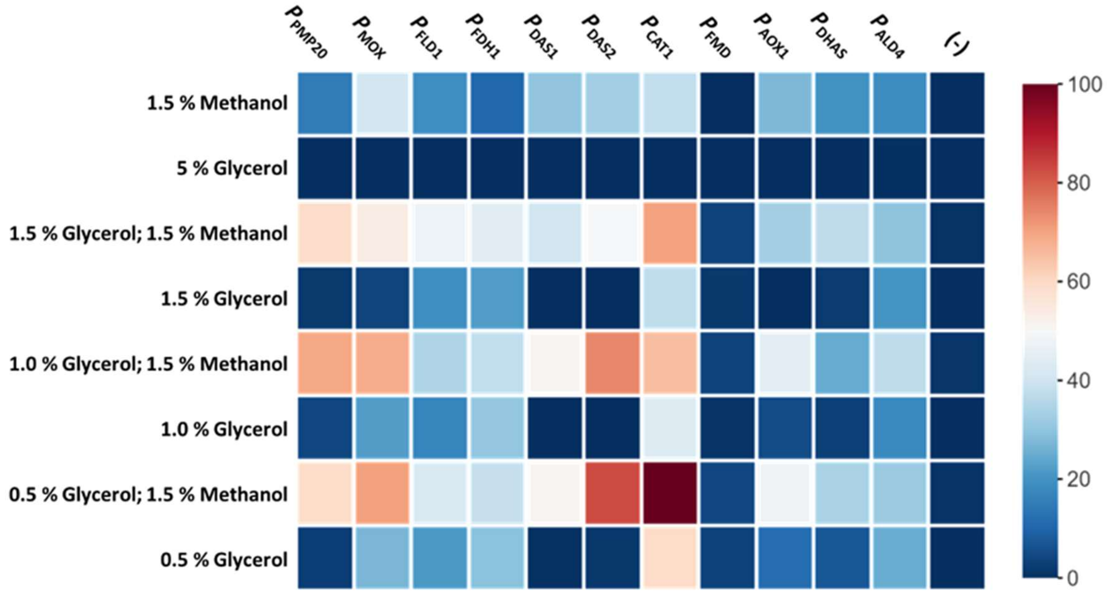

**Supplemental Figure 3 Time course of derepression screening *Tte*UPO constructs**

Respective constructs ($P_X$- *Sce*−Invertase 2 SP-*Tte*UPO and $P_{HpFMD}$- *Ani*−α Amylase SP-*Tte*UPO) were cultivated in 96 well plates for 100 h and individual samples retrieved at the indicated time points. For each time point two wells of the respective construct were unified, cells spun down and UPO activity in the supernatant (40 µL volume) determined by the means of NBD conversion measurements.

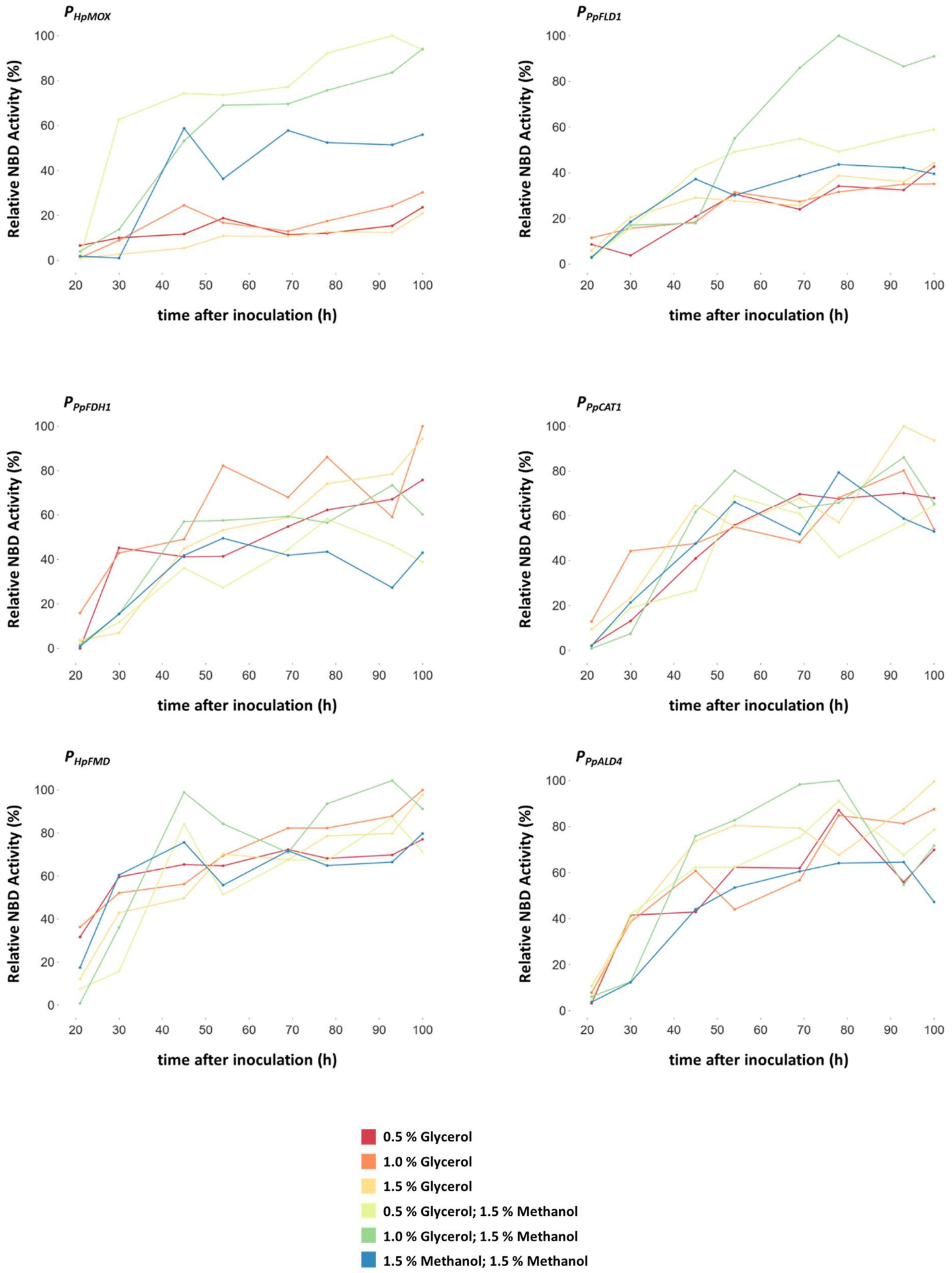

**Supplemental Figure 4 Testing synergistic effects of promoter and signal peptide usage**

Constructs of *Tte*UPO and the respective promoter/signal peptides were cultivated for 72 h (6 biological replicates) and UPO activity measured in 20 µL of supernatant. The highest mean activity was set as 100 % and all values normalized accordingly.

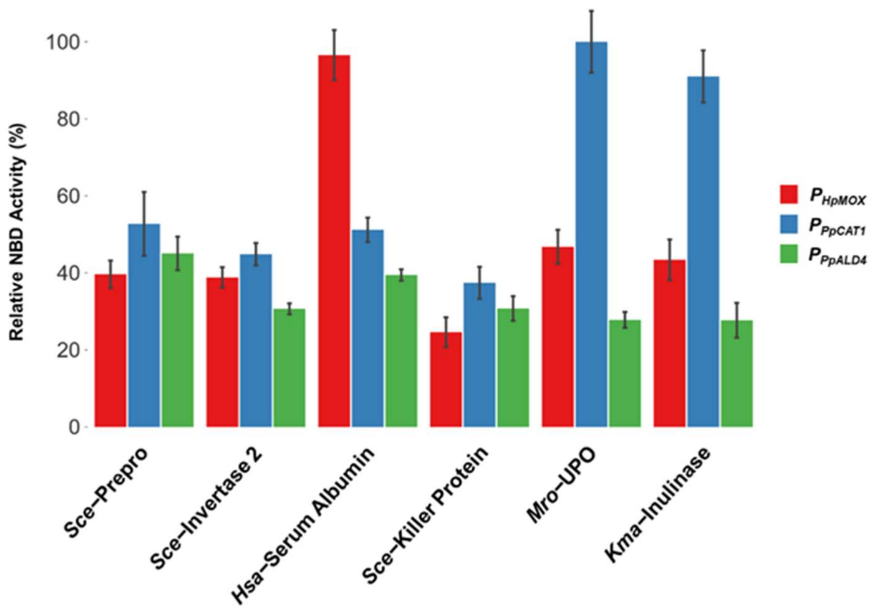


**Supplemental Figure 5 Screening landscape *Tte*UPO**

Primary screening results for the constructed promoter signal peptide shuffling library in combination with *Tte*UPO. 384 colonies (4 MTPs; 8 positive and negative controls respectively) were tested for NBD conversion. All clones have been ranked by their relative activity towards the substrate NBD. Included positive controls ($P_{PpCAT1}$-*Mro*–UPO) are indicated with a (+), the empty plasmid backbone (pPAP004) was used as negative control (-). 5 % of the activity of the positive controls was chosen as threshold for clones to be classified as active (dashed line). Out of all tested clones, in total 242 (66 %) exhibited clear activity towards NBD.

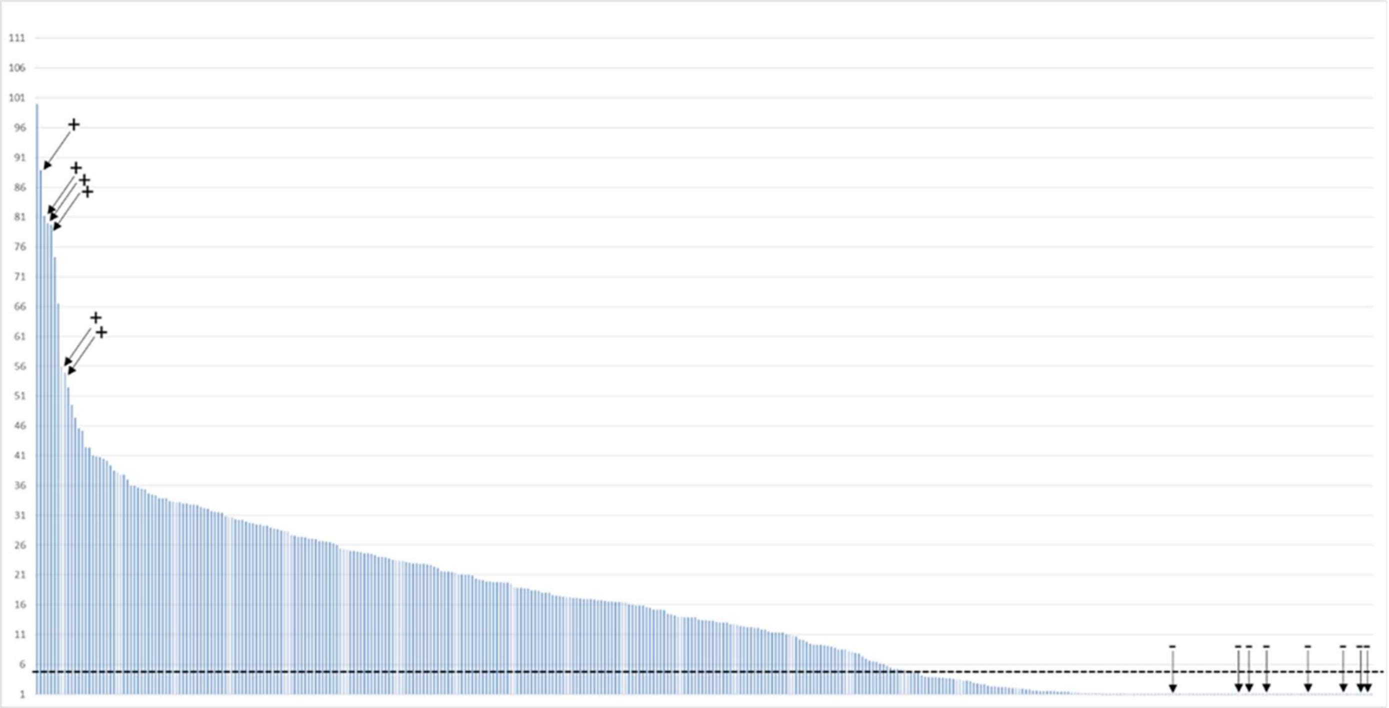

**Supplemental Figure 6 Screening landscape *Mth*UPO**

Primary screening results for the constructed promoter signal peptide shuffling library in combination with *Mth*UPO. 384 colonies (4 MTPs; 8 positive and negative controls respectively) were tested for NBD conversion. All clones have been ranked by their relative activity towards the substrate NBD. Included positive controls ($P_{Pp\mathrm{FLD1}}$- *Sce*−α Galactosidase) are indicated with a (+), the empty plasmid backbone (pPAP004) was used as negative control (-). 10 % of the activity of the positive controls was chosen as threshold for clones to be classified as active (dashed line). Out of all tested clones, in total 205 (56 %) exhibited clear activity towards NBD.

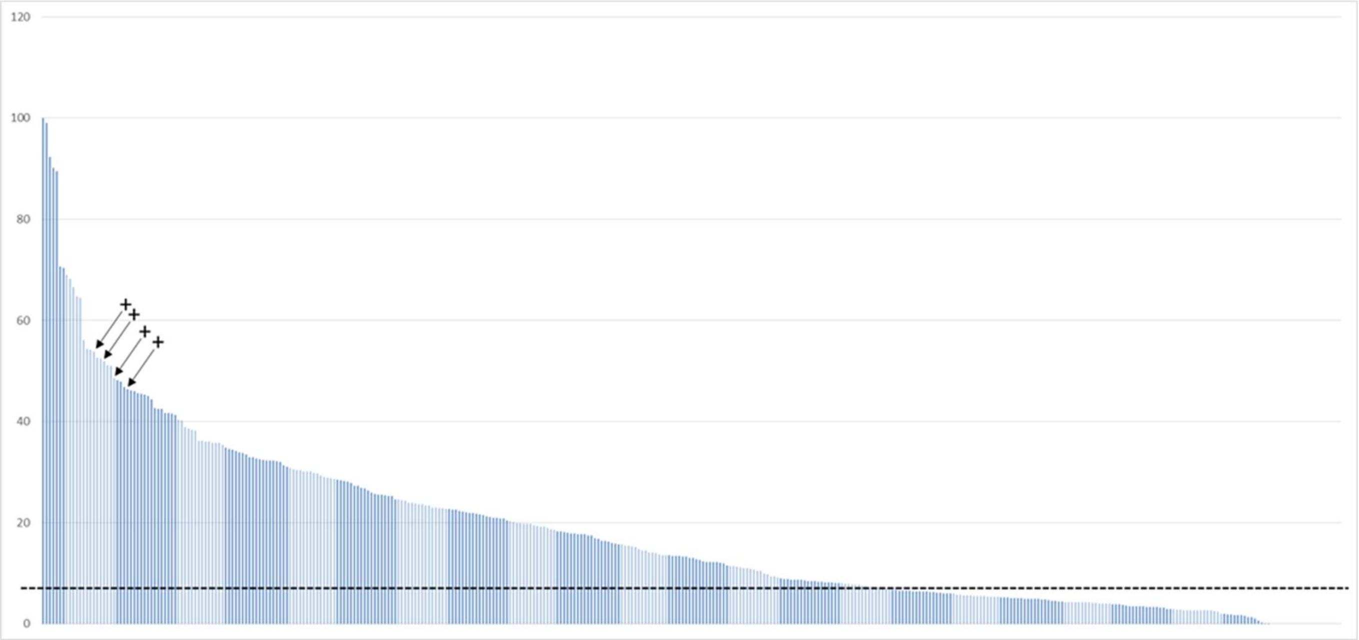


**Supplemental Figure 7 Screening landscape *Aae*UPO***

Primary screening results for the constructed promoter signal peptide shuffling library in combination with *Aae*UPO*. 384 colonies (4 MTPs; 8 negative controls respectively) were tested for NBD conversion. All clones have been ranked by their relative activity towards the substrate NBD. Included positive controls ($P_{Pp\mathrm{CAT1}}$- Gma-UPO) are indicated with a (+), the empty plasmid backbone (pPAP004) was used as negative control (-). 20 % of the activity of the positive controls was chosen as threshold for clones to be classified as active (dashed line). Out of all tested clones, in total 28 (7 %) exhibited clear activity towards NBD.

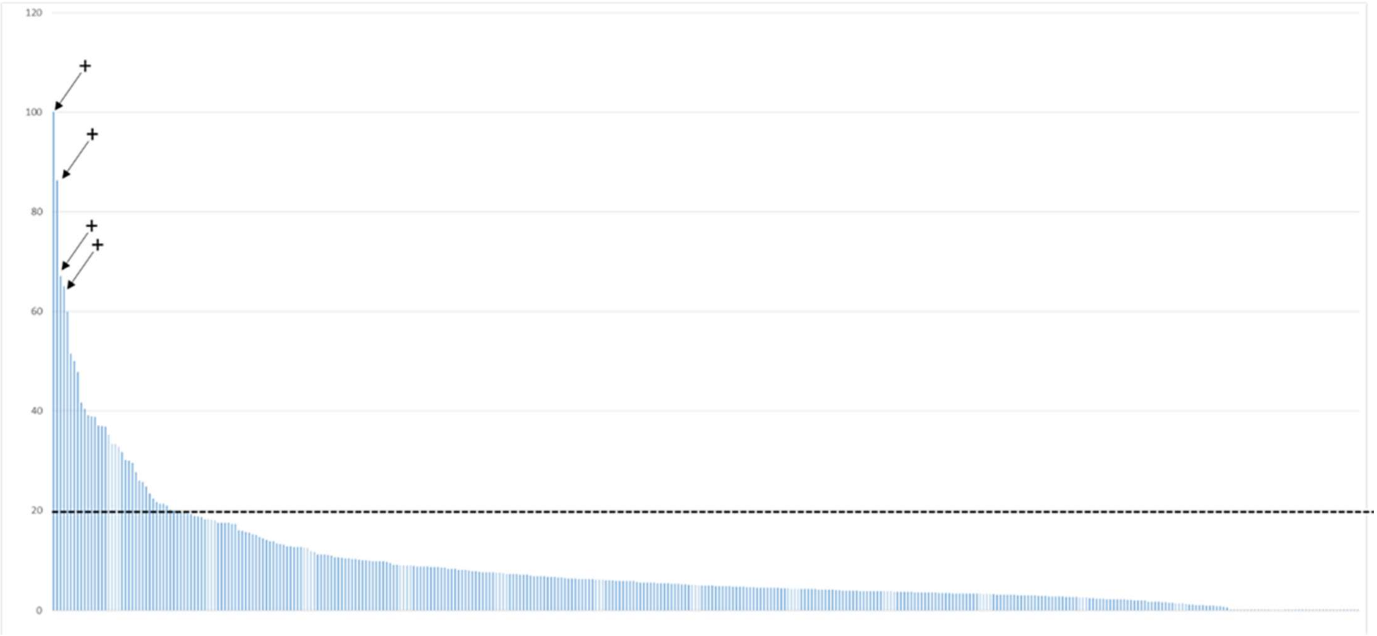

**Supplemental Figure 8 Activity distribution of carbon source screening episomal/integrative constructs *Tte*UPO, *Mth*UPO, *Aae*UPO***

6 biological replicates of the respective episomal and integrative constructs were tested towards NBD conversion. The highest sample mean within one group (out of the 12 tested conditions) was set as 100 % activity and all conversion values normalized accordingly.

**Scheme of Boxplots:**

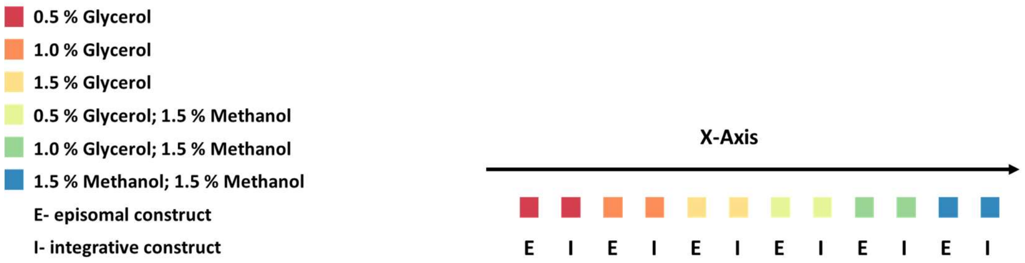


***Tte*UPO**

$P_{HpFMD}$- *Ani*–α Amylase

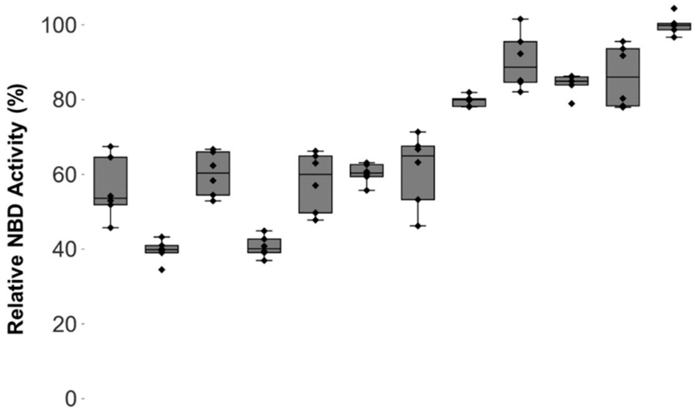


$P_{PpCAT1}$- *Mro*–UPO

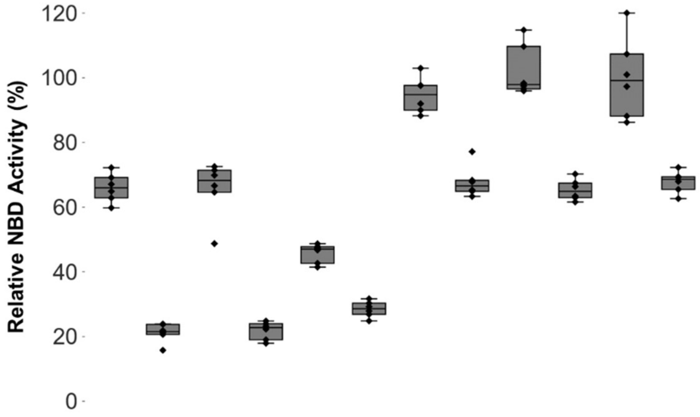

***Mth*UPO**

$P_{HpFMD}$- *Gga*–Lysozym

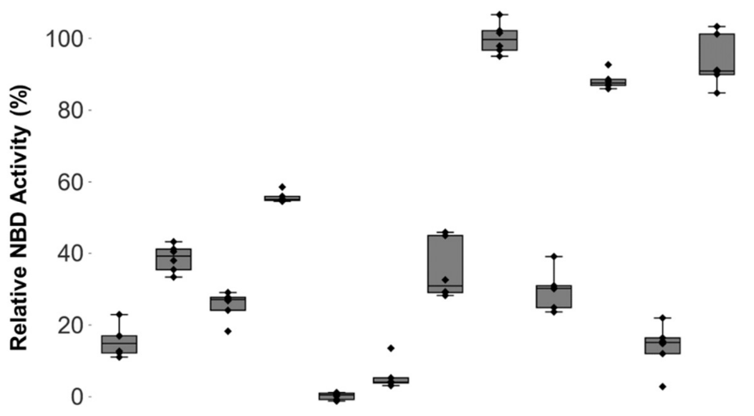


$P_{PpFLD1}$- *Hsa*–Serum Albumin

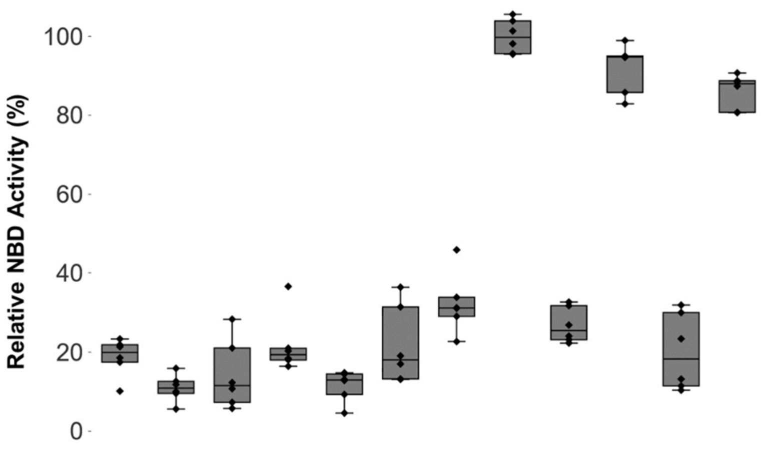

***Aae*UPO***

$P_{HpFMD}$- *Gma*–UPO

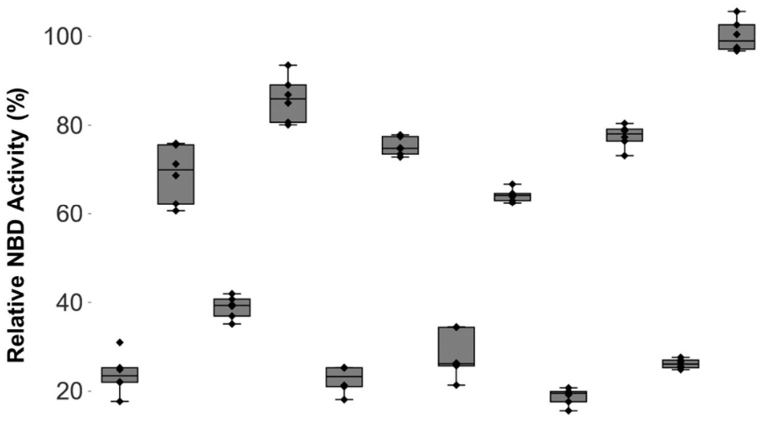


$P_{HpMOX}$- *Gma*–UPO

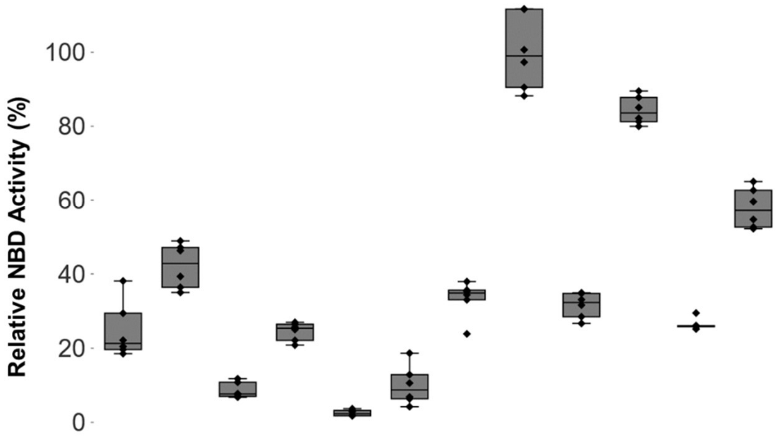

**Supplemental Figure 9 Screening landscape *Dca*UPO**

Primary screening results for the constructed promoter signal peptide shuffling library in combination with *Dca*UPO. 368 colonies (4 MTPs; 8 positive and negative controls respectively) were tested for DMP conversion. All clones have been ranked by their relative activity towards the substrate DMP. The empty plasmid backbone (pPAP004) was used as negative control (-). 35 % of the activity of the most active clones was chosen as threshold for clones to be classified as active (dashed line). Out of all tested clones, in total 96 (26 %) exhibited clear activity towards DMP.

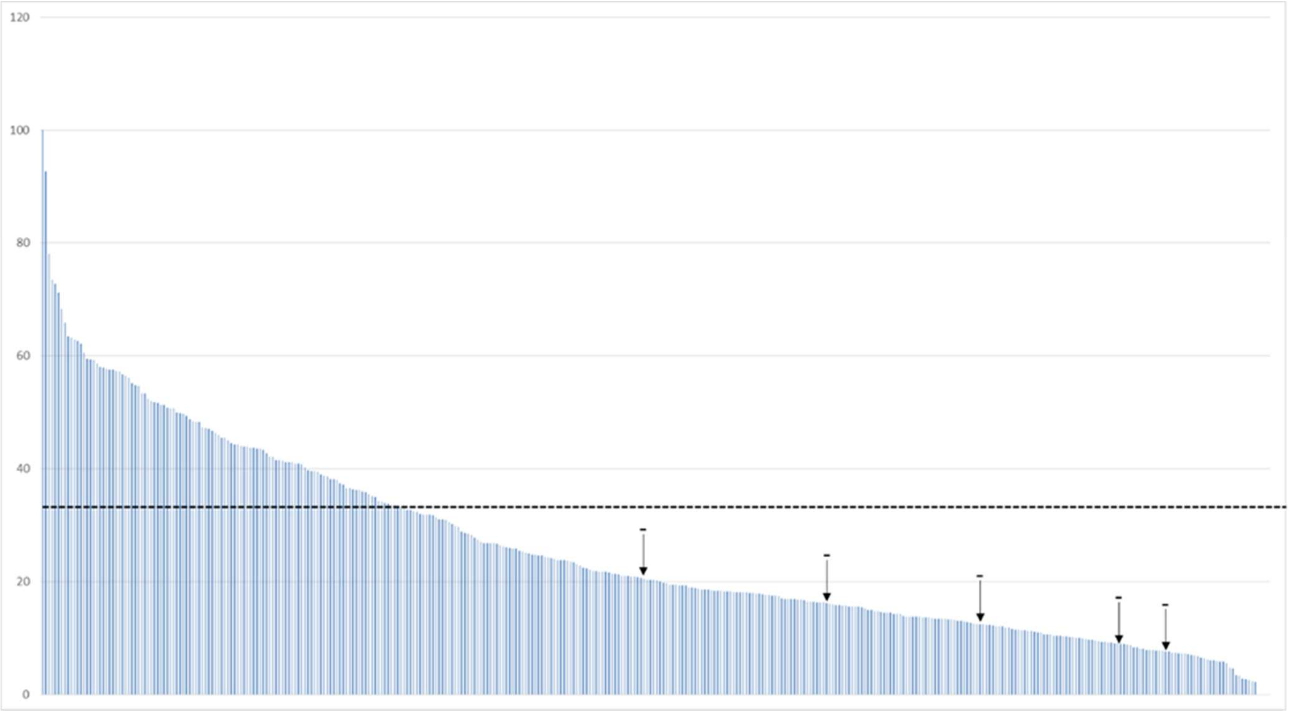

**Supplemental Figure 10 Screening landscape *Mfe*UPO**

Primary screening results for the constructed promoter signal peptide shuffling library in combination with *Mfe*UPO. 368 colonies (4 MTPs; 8 positive and negative controls respectively) were tested for NBD conversion. All clones have been ranked by their relative activity towards the substrate NBD. The empty plasmid backbone (pPAP004) was used as negative control (-). 15 % of the activity of the most active clones was chosen as threshold for clones to be classified as active (dashed line). Out of all tested clones, in total 89 (24 %) exhibited clear activity towards NBD.

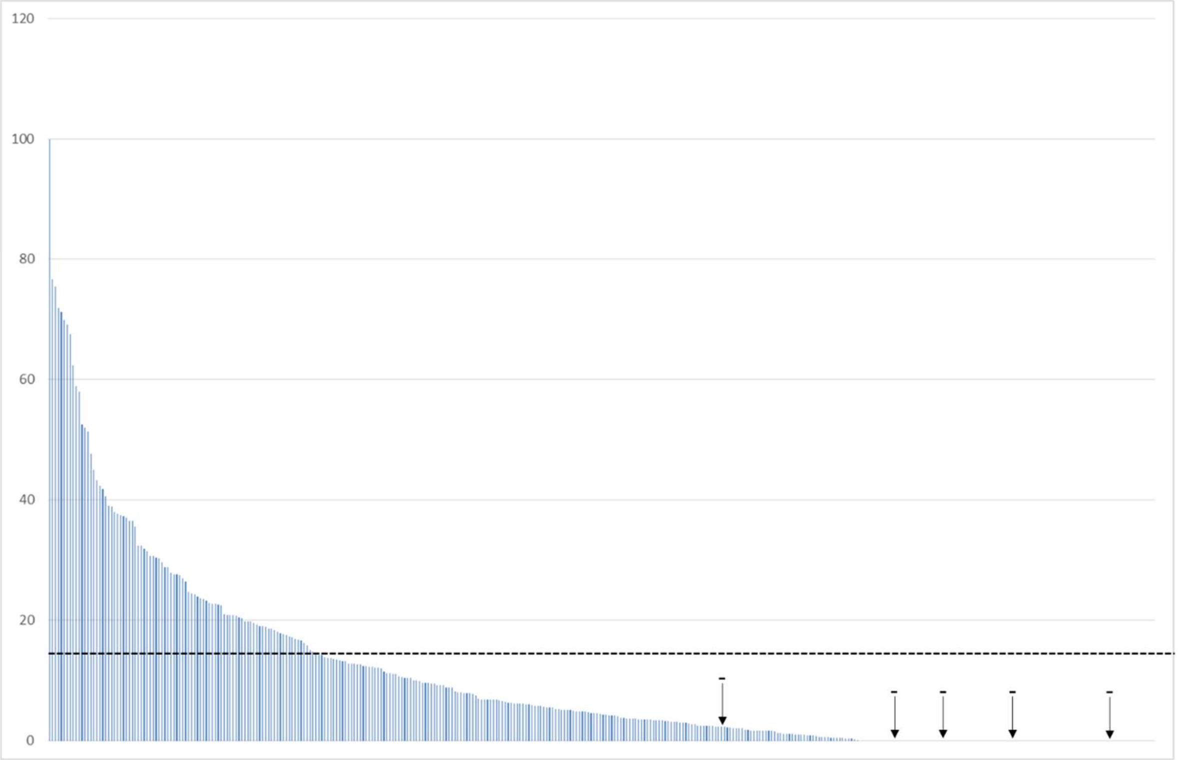

**Supplemental Figure 11 Screening landscape *Mhi*UPO**

Primary screening results for the constructed promoter signal peptide shuffling library in combination with *Mhi*UPO. 368 colonies (4 MTPs; 8 positive and negative controls respectively) were tested for NBD conversion. All clones have been ranked by their relative activity towards the substrate NBD. The empty plasmid backbone (pPAP004) was used as negative control (-) and the construct $P_{HpFMD}$-*Gga*-Lysozym SP-*Mth*UPO as positive control (+). 10 % of the activity of the most active clones was chosen as threshold for clones to be classified as active (dashed line). Out of all tested clones, in total 113 (31 %) exhibited clear activity towards NBD.

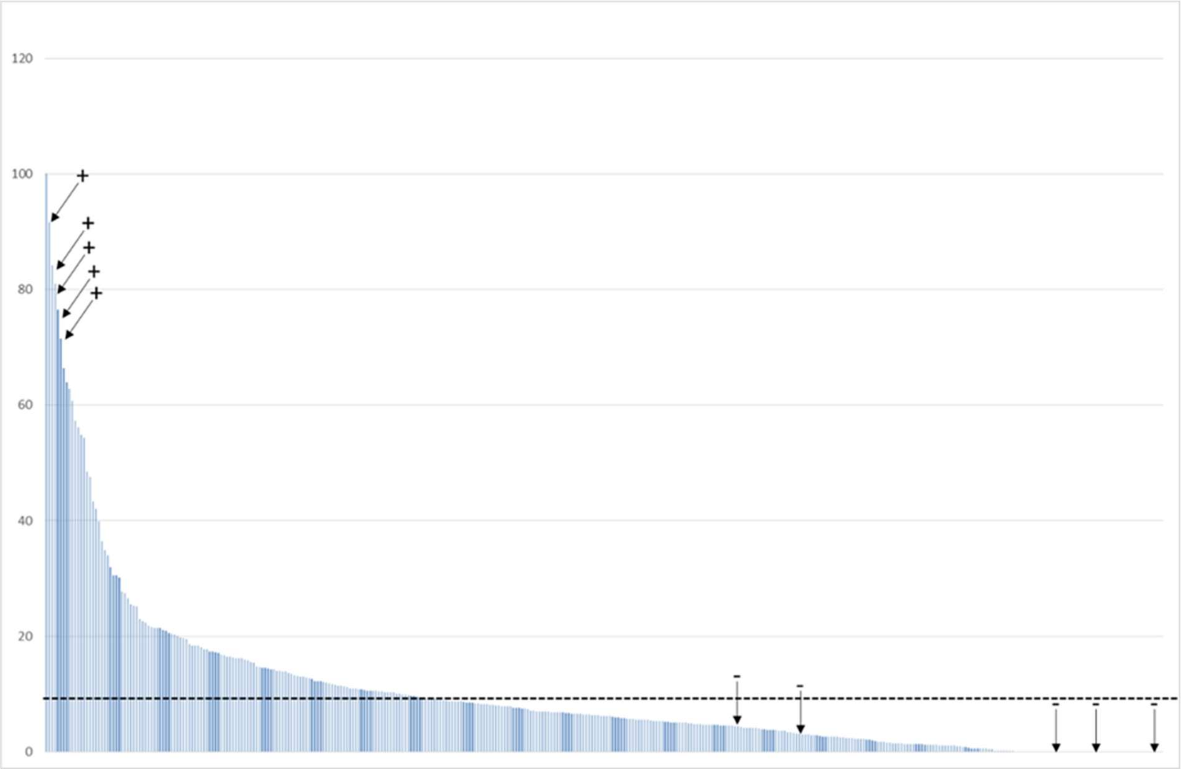

**Supplemental Figure 12 Activity distribution of carbon source screening episomal/integrative constructs *Mhi*UPO, *Mfe*UPO, *Dca*UPO**

6 biological replicates of the respective episomal and integrative constructs were tested towards DMP conversion. The highest sample mean within one group (out of the 12 tested conditions) was set as 100 % activity and all conversion values normalized accordingly.

**Scheme of Boxplots:**

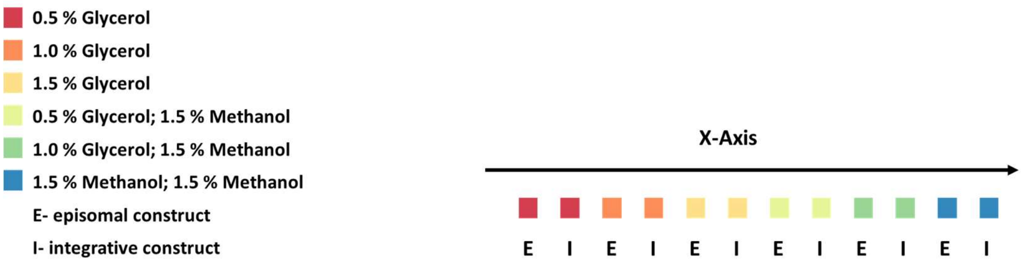


***Mhi*UPO**

$P_{PpALD4}$- *Aaw*–Glucoamylase

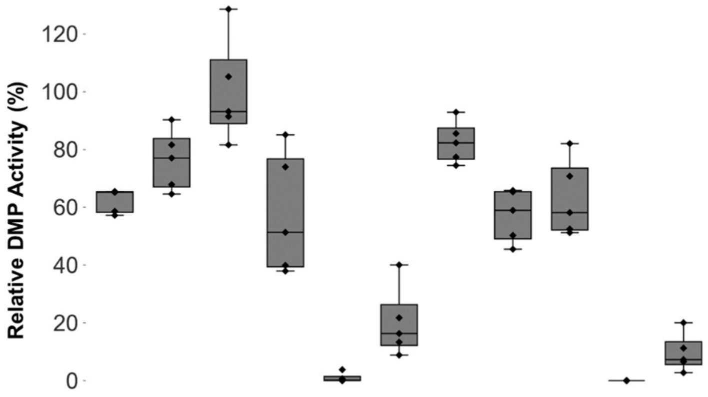


$P_{HpMOX}$- *Sce*–α Galactosidase

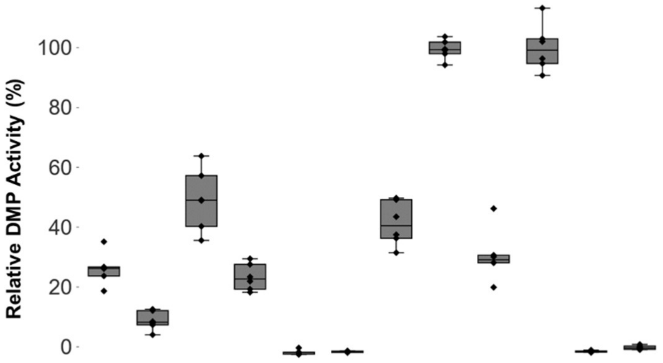

***Mfe*UPO**

$P_{HpFMD}$- *Gga*–Lysozym

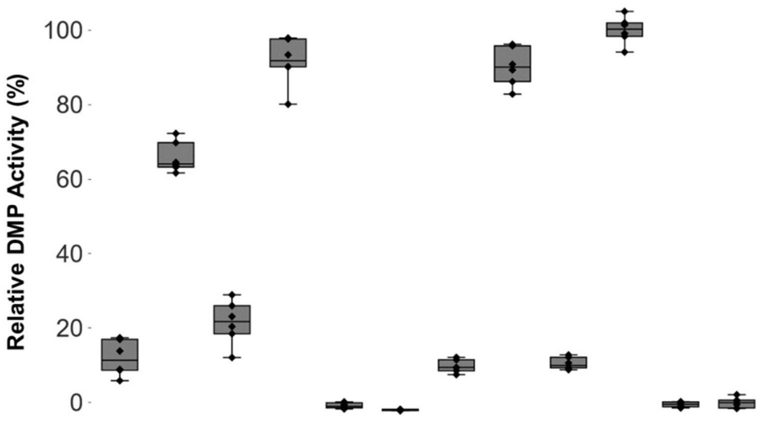


$P_{PpFLD1}$- *Ani*–α Amylase

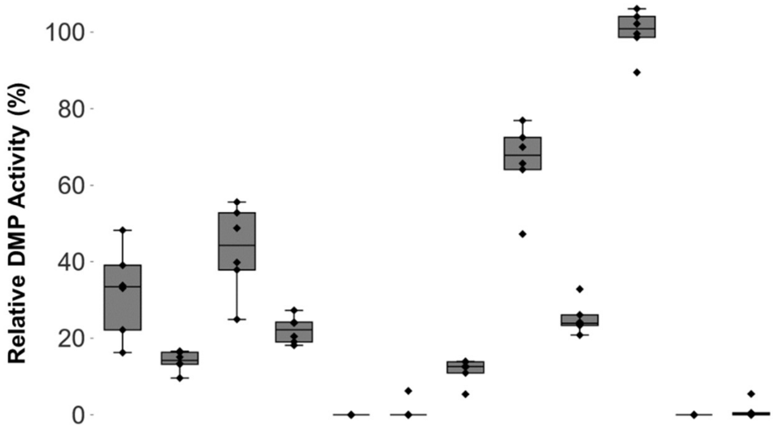

***Dca*UPO**

*P<sub>HpMOX</sub>*- *Sce*−α Galactosidase

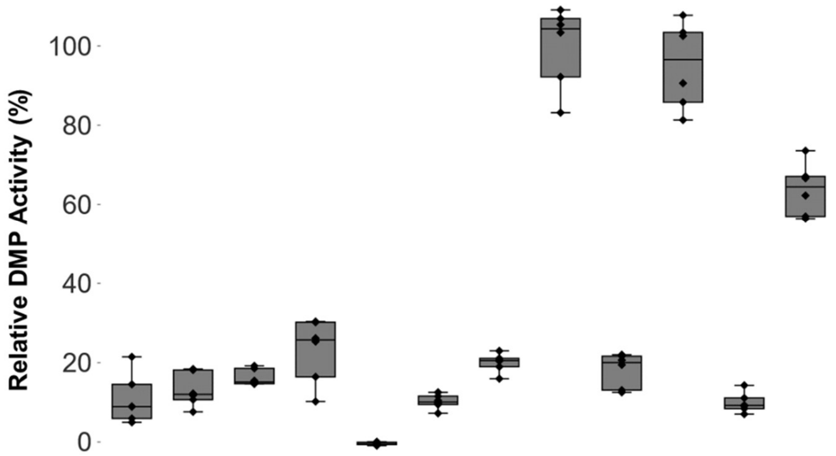


*P<sub>PpFLD1</sub>*- *Sce*−α Galactosidase

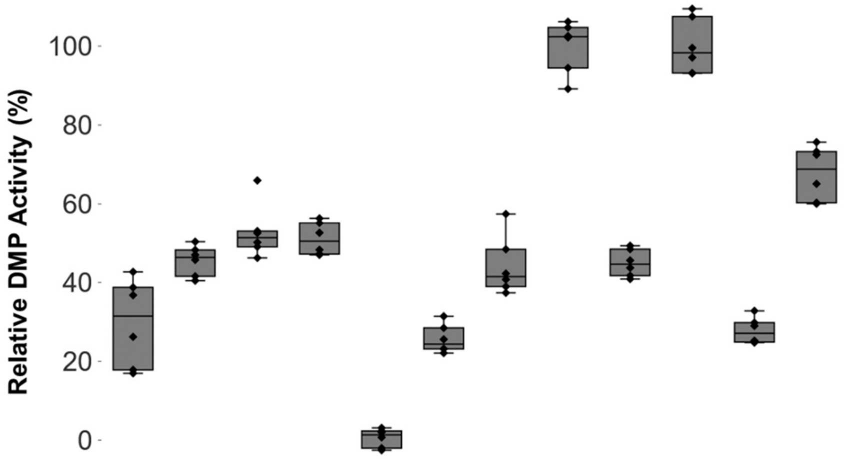

**Supplemental Figure 13 Absorption spectra UPOs and CO spectra**

Absorption spectra of the respective, purified UPOs were recorded in 100 mM potassium phosphate buffer (pH 7.0) in their native state (left). The reduced, CO spectrum (right) was recorded as differential spectrum after reduction with sodium dithionite and CO treatment using the reduced sample (without CO treatment) as blank measurement.

***Aae*UPO***

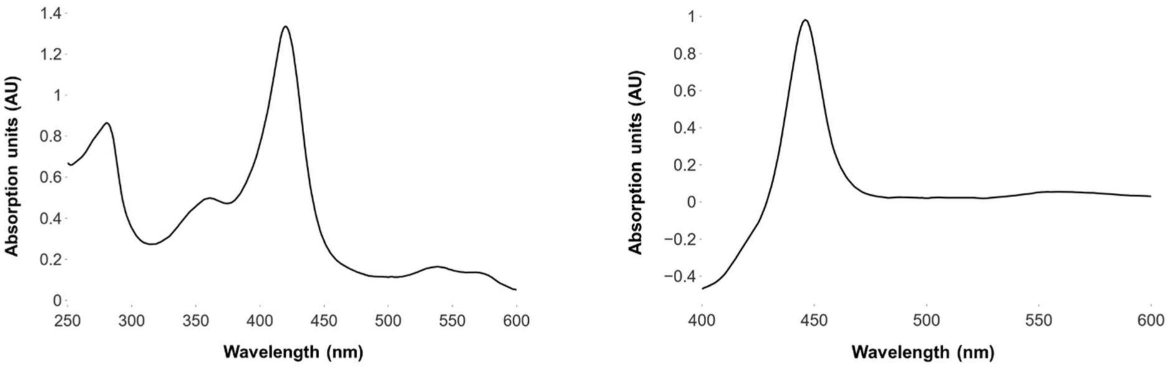


***Tte*UPO**

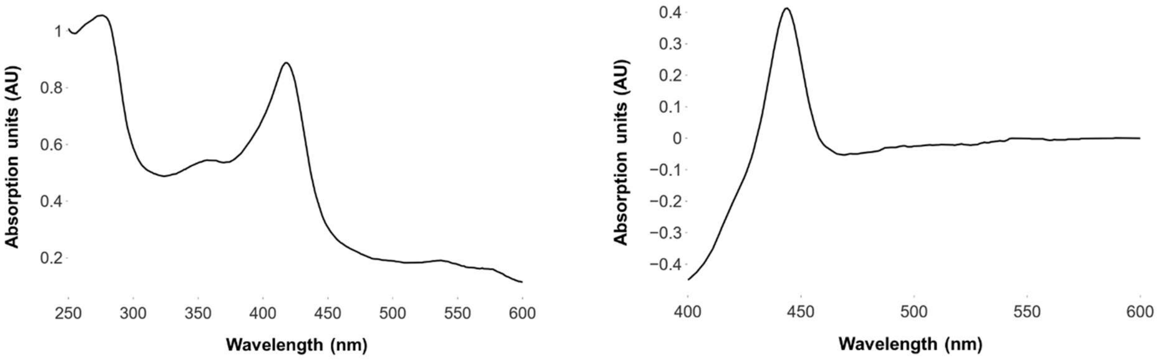


***Mth*UPO**

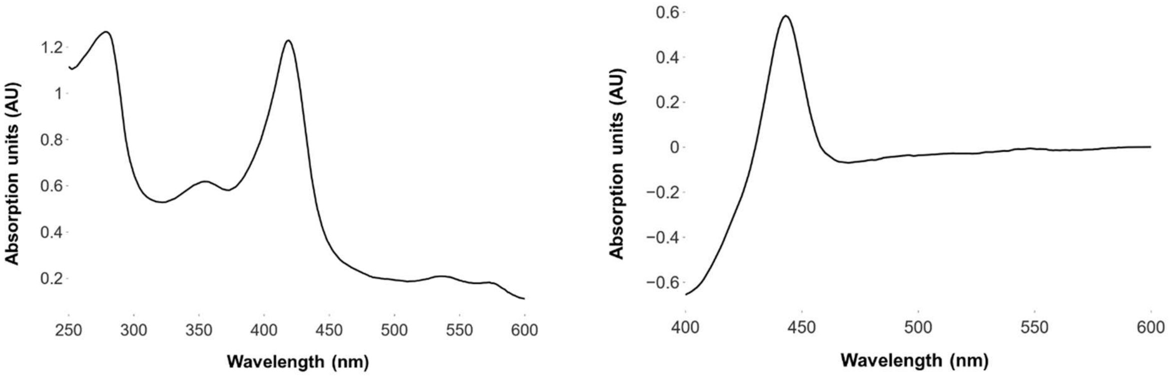

***Dca*UPO**

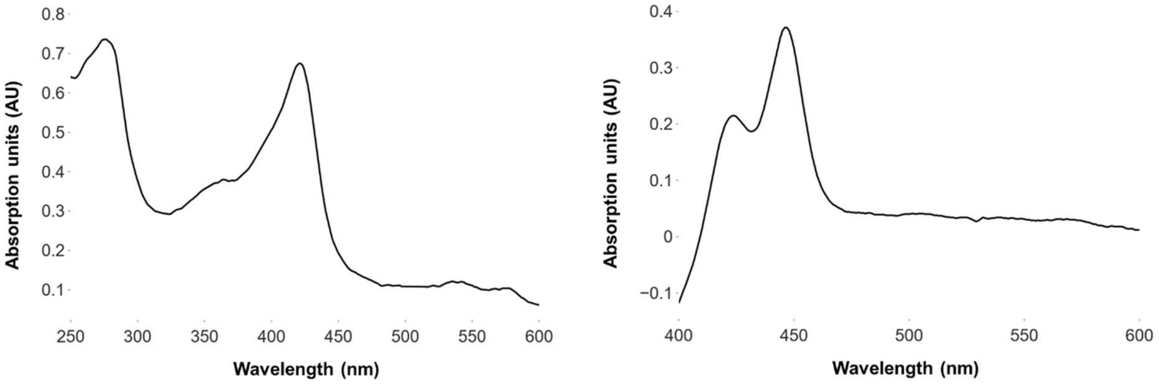


**Supplemental Figure 14 CO differential spectra of *Myceliophthora* UPOs**

Supernatant after cultivation was concentrated via ultrafiltration (approx. 10x fold concentrated) and 1 ml samples used for measurement (in 100 mM potassium phosphate; pH 6.0). Samples were reduced (+ sodium dithionite) and split into two halves- one half further treated with carbon monoxide. CO differential spectra were then recorded using the reduced, not CO treated sample as blank measurement.

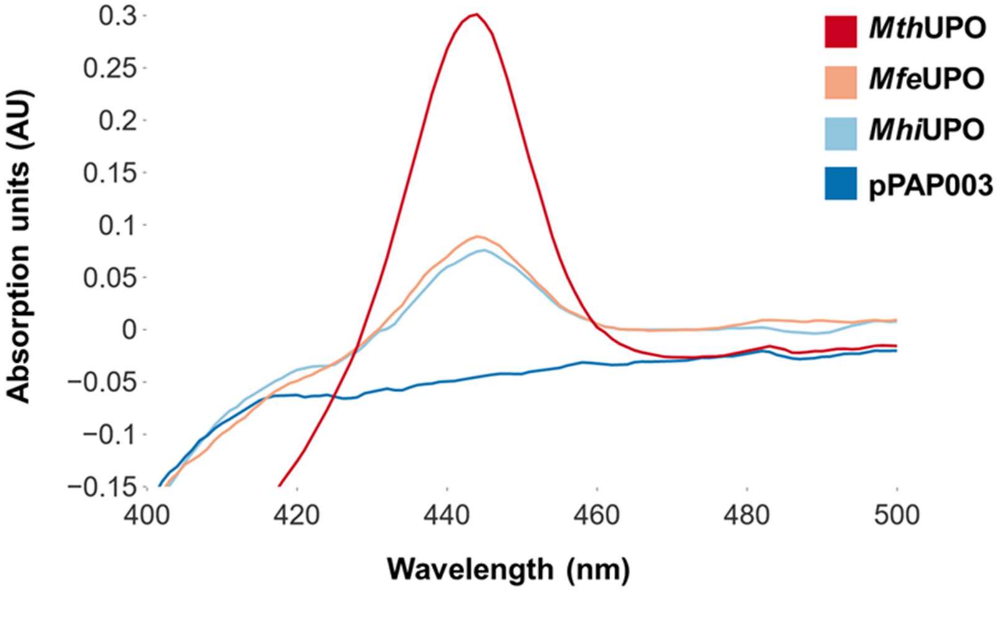

**Supplemental Figure 15 Screening landscape CalB**

Primary screening results for the constructed promoter signal peptide shuffling library in combination with CalB. 384 colonies (4 MTPs; 8 negative controls respectively) were tested for 4-Nitrophenyl Laurate conversion. All clones have been ranked by their relative activity towards the substrate. The empty plasmid backbone (pPAP004) was used as negative control (-). 20 % of the activity of the overall achieved activity was chosen as threshold for clones to be classified as active (dashed line). Out of all tested clones, in total 165 (44 %) exhibited clear, background distinguishable activity towards 4-Nitrophenyl laurate.

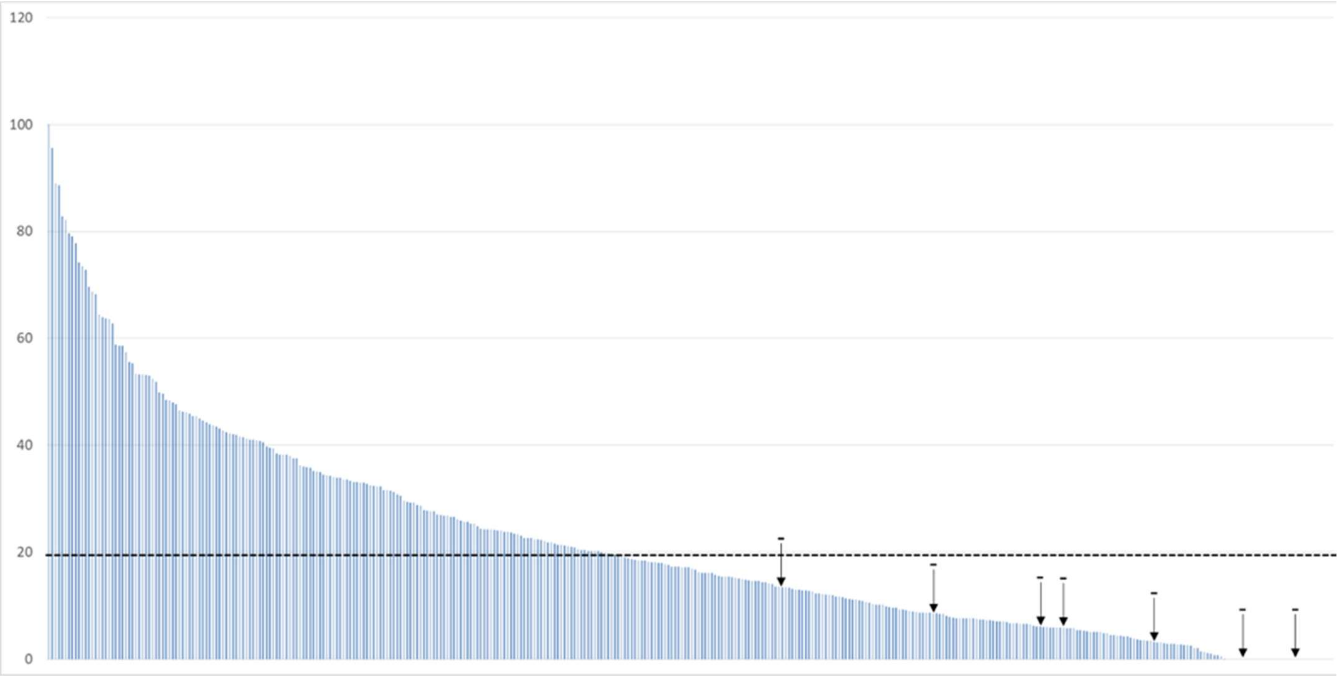

**Supplemental Figure 16 Screening landscape Mrl2**

Primary screening results for the constructed promoter signal peptide shuffling library in combination with Mrl2. 384 colonies (4 MTPs; 8 negative controls respectively) were tested for ABTS conversion. All clones have been ranked by their relative activity towards the substrate. The empty plasmid backbone (pPAP004) was used as negative control (-). 5 % of the activity of the overall achieved activity was chosen as threshold for clones to be classified as active (dashed line). Out of all tested clones, in total 263 (70 %) exhibited clear, background distinguishable activity towards ABTS.

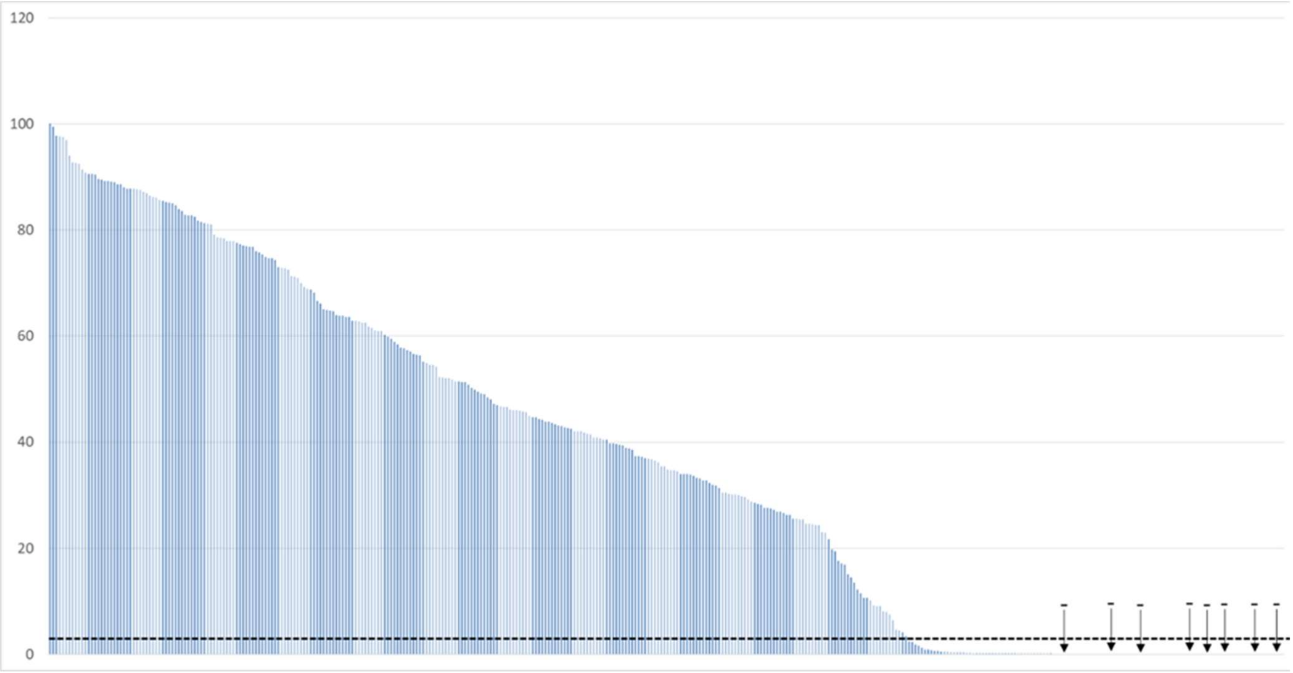

**Supplemental Figure 17 Activity distribution of carbon source screening episomal/integrative constructs CalB and Mrl2**

6 biological replicates of the respective episomal and integrative constructs were tested towards DMP (Mrl2) and 4-Nitrophenyl Laurate (CalB) conversion. The highest sample mean within one group (out of the 12 tested conditions) was set as 100 % activity and all conversion values normalized accordingly.

**Scheme of Boxplots:**

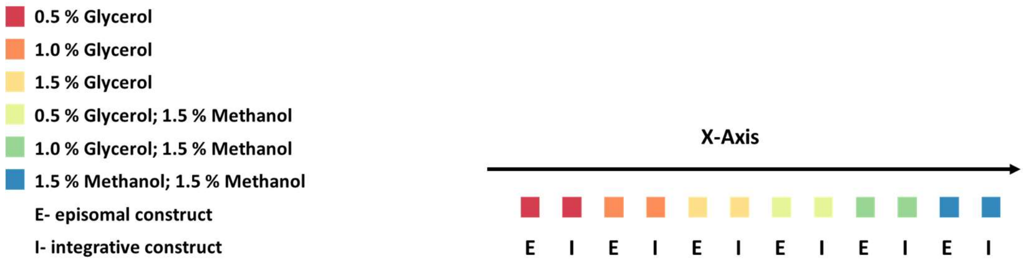


**CalB**

*$P_{HpFMD}$*- Sce–Killer Protein

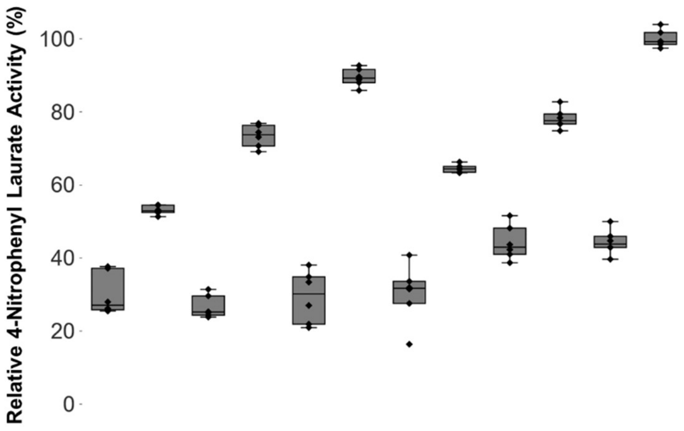


*$P_{PpFLD1}$*- *Ani*–α Amylase

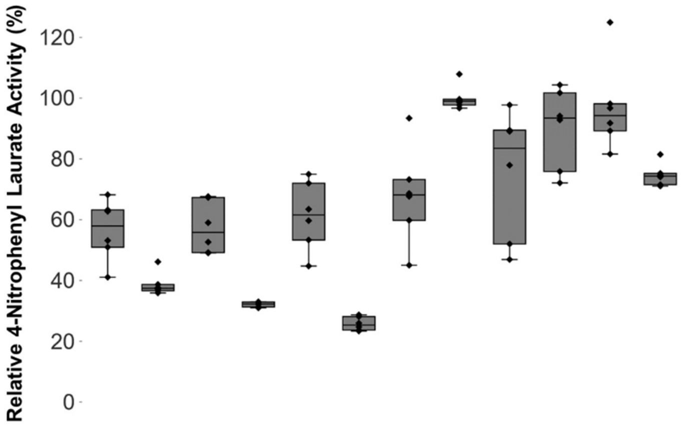

**Mrl2**

$P_{HpFMD}$- *Aaw*–Glucoamylase

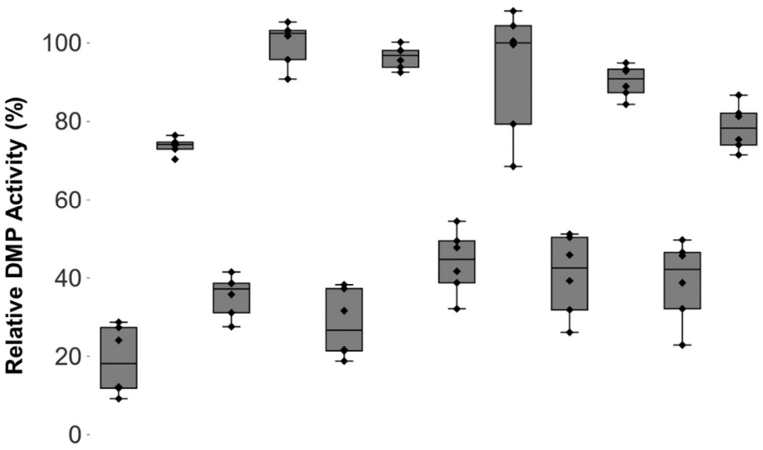


$P_{PpFLD1}$- *Cgl*–UPO

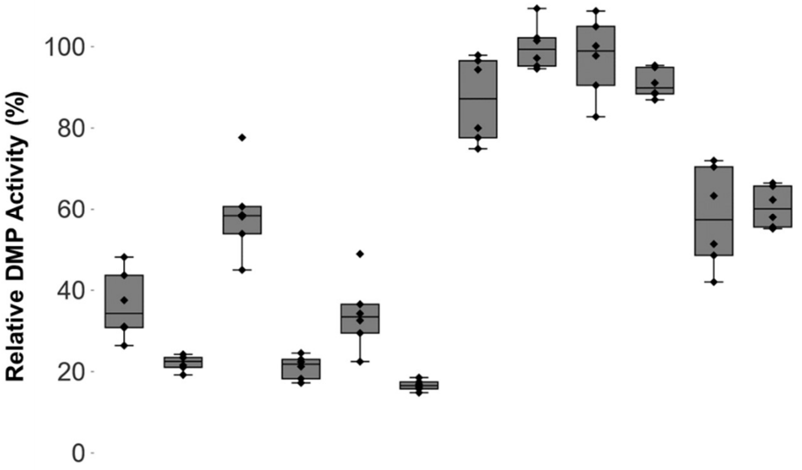

**Supplemental Table 1 Overview of oligonucleotides for sequencing of the created plasmids**

| Name | Sequence (5´ → 3´) |
|---|---|
| pAGM9121_for | CCTGTCGGGTTTCGCCACCT |
| pAGM9121_rev | GCCGTTACCACCGCTGCGTT |
| tGAP_rev | TCATTATGGCTGTATCTACTTTAGCGTA |
| pPAP004_for | GCAACGCGGCCTTTTTACGGTTCCTGCC |
| MthUPO_rev | GTATTCAACATCGGACAAGGAGCCCTAAC |
| TteUPO_rev | GTGTTCAGCATTGGACAAGGACCTCTACG |
| AaeUPO*_rev | CCACGAATATCGCCAGGTCGAAGCGGCTTCC |
| CalB_rev | CAGGTTAAACCGGCGTCAAGCACCGAC |
| Mr2Laccase_rev | GATCTTGTAAAACCATCCGGACTGACGTCTTGATTA |
| MfeUPO_rev | GCCATGATTGGCCAAGGTATTTAACATTGGACA |
| MhiUPO_rev | GGTTTGCCAAGGTGTTTAACATAGGACATGGAG |
| DcaUPO_rev | GCCAGAGTGTTCAACATAGGGCATGGTCC |

**Supplemental Table 2 Nucleotide Sequences of the MUT promoters (as cloned in pAGM9121)**

**Kozak Sequence**

**Golden Gate Assembly overhangs**

***$P_{PpPMP20}$***

AACGTTCTGGAGTGTCAAAACAGTAGTGATAAAAGGCTATGAAGGAGGTTGTCTAGGGGCTCGCGGAGGAAAGTGATTCA
AACAGACCTGCCAAAAAGAGAAAAAAGAGGGAATCCCTGTTCTTTCCAATGGAAATGACGTAACTTTAACTTGAAAAATAC
CCCAACCAGAAGGGTTCAAACTCAACAAGGATTGCGTAATTCCTACAAGTAGCTTAGAGCTGGGGGAGAGACAACTGAAG
GCAGCTTAACGATAACGCGGGGGGATTGGTGCACGACTCGAAAGGAGGTATCTTAGTCTTGTAACCTCTTTTTCCAGAGGC
TATTCAAGATTCATAGGCGATATCGATGTGGAGAAGGGTGAACAATATAAAAGGCTGGAGAGATGTCAATGAAGCAGCTG
GATAGATTTCAAATTTTCTAGATTTCAGAGTAATCGCACAAAACGAAGGAATCCCACCAAGCAAAAAAAAAAAATCTAAGAT
CATACAAAATG

***$P_{HpMOX}$***

AACGCGACGCGGAGAACGATCTCCTCGAGCTGCTCGCGGATCAGCTTGTGGCCCGGTAATGGAACCAGGCCGACGGCACGC
TCCTTGCGGACCACGGTGGCTGGCGAGCCCAGTTTGTGAACGAGGTCGTTTAGAACGTCCTGCGCAAAGTCCAGTGTCAGA
TGAATGTCCTCCTCGGACCAATTCAGCATGTTCTCGAGCAGCCATCTGTCTTTGGAGTAGAAGCGTAATCTCTGCTCCTCGTT
ACTGTACCGGAAGAGGTAGTTTGCCTCGCCGCCCATAATGAACAGGTTCTCTTTCTGGTGGCCTGTGAGCAGCGGGGACGTC
TGGACGGCGTCGATGAGGCCCTTGAGGCGCTCGTAGTACTTGTTCGCGTCGCTGTAGCCGGCCGCGGTGACGATACCCACA
TAGAGGTCCTTGGCCATTAGTTTGATGAGGTGGGGCAGGATGGGCGACTCGGCATCGAAATTTTTGCCGTCGTCGTACAGTG
TGATGTCACCATCGAATGTAATGAGCTGCAGCTTGCGATCTCGGATGGTTTTGGAATGGAAGAACCGCGACATCTCCAACA
GCTGGGCCGTGTTGAGAATGAGCCGGACGTCGTTGAACGAGGGGGCCACAAGCCGGCGTTTGCTGATGGCGCGGCGCTCGT
CCTCGATGTAGAAGGCCTTTTCCAGAGGCAGTCTCGTGAAGAAGCTGCCAACGCTCGGAACCAGCTGCACGAGCCGAGACA
ATTCGGGGGTGCCGGCTTTGGTCATTTCAATGTTGTCGTCGATGAGGAGTTCGAGGTCGTGGAAGATTTCCGCGTAGCGGCG
TTTTGCCTCAGAGTTTACCATGAGGTCGTCCACTGCAGAGATGCCGTTGCTCTTCACCGCGTACAGGACGAACGGCGTGGCC
AGCAGGCCCTTGATCCATTCTATGAGGCCATCTCGACGGTGTTCCTTGAGTGCGTACTCCACTCTGTAGCGACTGGACATCT
CGAGACTGGGCTTGCTGTGCTGGATGCACCAATTAATTGTTGCCGCATGCATCCTTGCACCGCAAGTTTTTAAAACCCACTC
GCTTTAGCCGTCGCGTAAAACTTGTGAATCTGGCAACTGAGGGGGTTCTGCAGCCGCAACCGAACTTTTCGCTTCGAGGACG
CAGCTGGATGGTGTCATGTGAGGCTCTGTTTGCTGGCGTAGCCTACAACGTGACCTTGCCTAACCGGACGGCGCTACCCACT
GCTGTCTGTGCCTGCTACCAGAAAATCACCAGAGCAGCAGAGGGCCGATGTGGCAACTGGTGGGGTGTCGGACAGGCTGTT
TCTCCACAGTGCAAATGCGGGTGAACCGGCCAGAAAGTAAATTCTTATGCTACCGTGCAGTGACTCCGACATCCCCAGTTTT
TGCCCTACTTGATCACAGATGGGGTCAGCGCTGCCGCTAAGTGTACCCAACCGTCCCCACACGGTCCATCTATAAATACTGC
TGCCAGTGCACGGTGGTGACATCAATCTAAAGTACAAAAACAAATCATACAAAATG

***$P_{PpFLD1}$***

AACGGCATGCAGGAATCTCTGGCACGGTGCTAATGGTAGTTATCCAACGGAGCTGAGGTAGTCGATATATCTGGATATGCC
GCCTATAGGATAAAAACAGGAGAGGGTGAACCTTGCTTATGGCTACTAGATTGTTCTTGTACTCTGAATTCTCATTATGGGA
AACTAAACTAATCTCATCTGTGTGTTGCAGTACTATTGAATCGTTGTAGTATCTACCTGGAGGGCATTCCATGAATTAGTGA
GATAACAGAGTTGGGTAACTAGAGAGAATAATAGACGTATGCATGATTACTACACAACGGATGTCGCACTCTTTCCTTAGT
TAAAACTATCATCCAATCACAAGATGCGGGCTGGAAAGACTTGCTCCCGAAGGATAATCTTCTGCTTCTATCTCCCTTCCTC
ATATGGTTTCGCAGGGCTCATGCCCCTTCTTCCTTCGAACTGCCCGATGAGGAAGTCCTTAGCCTATCAAAGAATTCGGGAC
CATCATCGATTTTTAGAGCCTTACCTGATCGCAATCAGGATTTCACTACTCATATAAATACATCGCTCAAAGCTCCAACTTTG
CTTGTTCATACAATTCTTGATATTCACATCATACAAAATG

***$P_{PpFDH1}$***

AACGAAATGGCAGAAGGATCAGCCTGGACGAAGCAACCAGTTCCAACTGCTAAGTAAAGAAGATGCTAGACGAAGGAGAC
TTCAGAGGTGAAAAGTTTGCAAGAAGAGAGCTGCGGGAAATAAATTTTCAATTTAAGGACTTGAGTGCGTCCATATTCGTG
TACGTGTCCAACTGTTTTCCATTACCTAAGAAAAACATAAAGATTAAAAAGATAAACCCAATCGGGAAACTTTAGCGTGCC
GTTTCGGATTCCGAAAAACTTTTGGAGCGCCAGATGACTATGGAAAGAGGAGTGTACCAAAATGGCAAGTCGGGGGCTACT
CACCGGATAGCCAATACATTCTCTAGGAACCAGGGATGAATCCAGGTTTTTGTTGTCACGGTAGGTCAAGCATTCACTTCTT
AGGAATATCTCGTTGAAAGCTACTTGAAATCCCATTGGGTGCGGAACCAGCTTCTAATTAAATAGTTCGATGATGTTCTCTA
AGTGGGACTCTACGGCTCAAACTTCTACACAGCATCATCTTAGTAGTCCCTTCCCAAAACACCATTCTAGGTTTCGGAACGT
AACGAAACAATGTTCCTCTCTTCACATTGGGCCGTTACTCTAGCCTTCCGAAGAACCAATAAAAGGGACCGGCTGAAACGG
GTGTGGAAACTCCTGTCCAGTTTATGGCAAAGGCTACAGAAATCCCAATCTTGTCGGGATGTTGCTCCTCCCAAACGCCATA
TTGTACTGCAGTTGGTGCGCATTTTAGGGAAAATTTACCCCAGATGTCCTGATTTTCGAGGGCTACCCCCAACTCCCTGTGCT
TATACTTAGTCTAATTCTATTCAGTGTGCTGACCTACACGTAATGATGTCGTAACCCAGTTAAATGGCCGAAAAACTATTTA
AGTAAGTTTATTTCTCCTCCAGATGAGACTCTCCTTCTTTTCTCCGCTAGTTATCAAACTATAAACCTATTTTACCTCAAATA
CCTCCAACATCACCCACTTAAACATCATACAAAATG

***$P_{PpDAS1}$***

AACGAATGATATAAACAACAATTGAGTGACAGGTCTACTTTGTTCTCAAAAGGCCATAACCATCTGTTTGCATCTCTTATCA
CCACACCATCCTCCTCATCTGGCCTTCAATTGTGGGGAACAACTAGCATCCCAACACCAGACTAACTCCACCCAGATGAAAC
CAGTTGTCGCTTACCAGTCAATGAATGTTGAGCTAACGTTCCTTGAAACTCGAATGATCCCAGCCTTGCTGCGTATCATCCC
TCCGCTATTCCGCCGCTTGCTCCAACCATGTTTCCGCCTTTTTCGAACAAGTTCAAATACCTATCTTTGGCAGGACTTTTCCT
CCTGCCTTTTTTAGCCTCAGTTCTCGGTTAGCCTCTAGGCAAATTCTGTTCTTCATACCTATATCAACTTTTCATCAGATAGCC
TTTGGGTTCAAAAAAGAACTAAAGCAGGATGCCTGATATATAAATCCCAGATGATCTGCTTTTGAAACTATTTTCAGTATCT
TGATTCGTTTACTTACAAACAACTATTGTTGATTTTATCTGGAGAATAATCGAACATCATACAAAATG

***$P_{PpDAS2}$***

AACGCAATCTGTGGTTTGCTAAACTGGAAGTCTGGTAAGGACTCTAGCAAGTCCGTTACTCAAAAAGTCATACCAAGTAAG
ATTACGTAACACCTGGGCATGACTTTCTAAGTTAGCAAGTCACCAAGAGGGTCCTATTTAACGTTTGGCGGTATCTGAAACA
CAAGACTTGCCTATCCCATAGTACATCATATTACCTGTCAAGCTATGCTACCCCACAGAAATACCCCAAAAGTTGAAGTGAA
AAAATGAAAATTACTGGTAACTTCACCCCATAACAAACTTAATAATTTCTGTAGCCAATGAAAGTAAACCCCATTCAATGTT
CCGAGATTTAGTATACTTGCCCCTATAAGAAACGAAGGATTTCAGCTTCCTTACCCCATGAACAGAAATCTTCCATTTACCC
CCCACTGGAGAGATCCGCCCAAACGAACAGATAATAGAAAAAAGAAATTCGGACAAATAGAACACTTTCTCAGCCAATTA
AAGTCATTCCATGCACTCCCTTTAGCTGCCGTTCCATCCCTTTGTTGAGCAACACCATCGTTAGCCAGTACGAAAGAGGAAA
CTTAACCGATACCTTGGAGAAATCTAAGGCGCGAATGAGTTTAGCCTAGATATCCTTAGTGAAGGGTTGTTCCGATACTTCT
CCACATTCAGTCATAGATGGGCAGCTTTGTTATCATGAAGAGACGGAAACGGGCATTAAGGGTTAACCGCCAAATTATATA
AAGACAACATGTCCCCAGTTTAAAGTTTTTCTTTCCTATTCTTGTATCCTGAGTGACCGTTGTGTTTAATATAACAAGTTCGT
TTTAACTTAAGACCAAAACCAGTTACAACAAATTATAACCCCTCTAAACACTAAAGTTCACTCTTATCAAACTATCAAACAT
CAAAATCATACAAAATG

***$P_{PpCAT1}$***

AACGTAATCGAACTCCGAATGCGGTTCTCCTGTAACCTTAATTGTAGCATAGATCACTTAAATAAACTCATGGCCTGACATC
TGTACACGTTCTTATTGGTCTTTTAGCAATCTTGAAGTCTTTCTATTGTTCCGGTCGGCATTACCTAATAAATTCGAATCGAG
ATTGCTAGTACCTGATATCATATGAAGTAATCATCACATGCAAGTTCCATGATACCCTCTACTAATGGAATTGAACAAAGTT
TAAGCTTCTCGCACGAGTCCGAATCCATACTATGCACCCCTCAAAGTTGGGATTAGTCAGGAAAGCTGAGCAATTAACTTCC
CTCGATTGGCCTGGACTTTTCGCTTAGCCTGCCGCAATCGGTAAGTTTCATTATCCCAGCGGGGTGATAGCCTCTGTTGCTCA
TCAGGCCAAAATCATATATAAGCTGTAGACCCAGCACTTCAATTACTTGAAATTCACCATAACACTTGCTCTAGTCAAGACT
TACAATTAAATCATACAAAATG

***$P_{PpAOX1}$***

AACGAGACGAAAGGTTGAATGAAACCTTTTTGCCATCCGACATCCACAGGTCCATTCTCACACATAAGTGCCAAACGCAAC
AGGAGGGGATACACTAGCAGCAGACCGTTGCAAACGCAGGACCTCCACTCCTCTTCTCCTCAACACCCACTTTTGCCATCGA
AAAACCAGCCCAGTTATTGGGCTTGATTGGAGCTCGCTCATTCCAATTCCTTCTACTAGGCTACTAACACCGTGACTTTATTA
GCCTGTCTATCCTGGCCCCCCTGGCGAGGTTCATGTTTGTTTATTTCCGAATGCAACAAGCTCCGCATTACACCCGAACATC
ACTCCAGATGAGGGCTTTCTGAGTGTGGGGTCAAATAGTTTCATGTTCCCCAAATGGCCCAAAACTGACAGTTTAAACGCTG
TCTTGGAACCTAATATGACAAAAGCGTGATCTCATCCAAGATGAACTAAGTTTGGTTCGTTGAAATGCTAACGGCCAGTTGG
TCAAAAAGAAACTTCCAAAAGTCGGCATACCGTTTGTCTTGTTTGGTATTGATTGACGAATGCTCAAAAATAATCTCATTAAT
GCTTAGCGCAGTCTCTCTATCGCTTCTGAACCCCGGTGCACCTGTGCCGAAACGCAAATGGGGAAACACCCGCTTTTTGGAT
GATTATGCATTGTCTCCACATTGTATGCTTCCAAGATTCTGGTGGGAATACTGCTGATAGCCTAACGTTCATGATCAAAATTT
AACTGTTCTAACCCCTACTTGACAGCAATATATAAACAGAAGGAAGCTGCCCTGTCTTAAACCTTTTTTTTTATCATCATTAT
TAGCTTACTTTCATAATTGCGACTGGTTCCAATTGACAAGCTTTTGATTTTAACGACTTTTAACGACAACTTGAGAAGATCA
AAAAACAACTAATTATTCGAAACATCATACAAAATG

***$P_{PpALD4}$***

AACGGACAATAAGAAGAAATAAAAAGAAAAGCGGTGGGGGAGGGATTATTAAATAAGGATTATGTAACCCCAGGGTACCG
TTCTATACATATTTAAGGATTATTTAGGACAATCGATGAAATCGGCATCAAACTGGATGGGAGTATAGTGTCCGGATAATCG
GATAAATCATCTTGCGAGGAGCCGCTTGGTTGGTTGGTGAGAGGAGTGAAATATGTGTCTCCTCACCCAAGAATCGCGATA
TCAGCACCCTGTGGGGGACACTATTGGCCTCCCTCCCAAACCTTCGATGTGGTAGTGCTTTATTATATTGATTACATTGATTA
CATAGCTAAACCCTGCCTGGTTGCAAGTTGAGCTCCGAATTCCAATATTAGTAAAATGCCTGCAAGATAACCTCGGTATGGC
GTCCGACCCCGCTTAATTATTTTAACTCCTTTCCAACGAGGACTTCGTAATTTTTGATTAGGGAGTTGAGAAACGGGGGGTC
TTGATACCTCCTCGATTTCAGATCCCACCCCCTCTCAGTCCCAAGTGGGACCCCCCTCGGCCGTGAAATGCGCGCACTTTAG
TTTTTTTCGCATGTAAACGCCGGTGTCCGTCAATTAAAAGTCGCAGACTAGGGTGAACTTTACCATTTTTGTCGCACTCCGTC
TCCTCGGAATAGGGGTGTAGTAATTCTGCAGTAGTGCAATTTTTACCCCGCCAAGGGGGGGCGAAAAGAGACGACCTCATC
ACGCATTCTCCAGTCGCTCTCTACGCCTACAGCACCGACGTAGTTAACTTTCTCCCATATATAAAGCAATTGCCATTCCCCTG
AAAACTTTAACCTCTGCTTTTTCTTGATTTTTCCTTGCCCAAAGAAAAGATCATACAAAATG

***$P_{HpDHAS}$***

AACGCTCGAGAGCCGTGGAACAGAGTCCCCTGAATCTCTTGCCAAGCGGCTTGCTGCTGCATCTGCGGAGATGGAGTACGC
CAGGGCAGTGGACACGACAAGGTCATTGTCAACGATGACCTTGAGAAGGCGTACTCTGAGCTGAAGGAGTTCATTTTCGCC
GAGCCCATCTAAGCATTCATAAATTTTTAATATCTAGAGCTCTCATACGGGACAGTATCTCCTCCAACCTTGCGTCAAGCTT
GTCCTCTTCATGCTCCTCAACAGTCATGGCATCCAGCTGCTGCTGCTTTTGCTCCAGCCTGGCATATATGTCGCCATACAGCT
TGAGTTGGATTTTGATGAAACTCTCAAAGGTAGGGTCCACCAGTGACAGTCGCAGCGCAATGAACTGCTCGATTTCGTTCTT
GAGCCGTGTGTTGATGTCCGTGTAGATATTTTCTGCCTCGTCGTACTCAACTTTGAACTTCTGCAGCTTGTCCAGGCTCTTCT
GTAACTGGTCTGTTTTCTCGGTGTGATGCTGCTCGGTCACCTGTCGCTCAATCGCTTCGTACTCGCTCTGCAGCTTCGAAACC
TTGAATCGTGAAACGTCGTAATCCACCTTTTTGCGTGCGCGCTTCTTGATCAGCTTGTTGATCTCGTCGTTGTACTTCTTCAG
CTCGTTAATCGGCTCCACGACCGTGATGCTCATTGGCTCCAGAATTTCTGGCAGAATATTGTCTTTGATGTCTACCACCATCT
GCAGATAATTCAGAGAAATACCATCTCTGGGGTTCACCTTGTGCTCTTCTGGCCGTTCCGCAGCTTCCGACCGCTTATCAGC
CTTGAGCTCAAAGCTATAGTCTCCGTAAAACGAGTCCAGTGTTCTAGCCATATTTATCTGAGTCTCGAGCAGATTCTCCGAA
ATTGCCCACAAAACGGCCTAGTTCCTGGTCCAGCTCGTTGGTGTAAGTCTCGAGTTTGCGGAAATTGGCCTCCTGGACGTCA
AACTCAGGATCAACAGAGGGCTCACCTTTGTTTGTGCGTAGTATCACATGTGCTCCGGCACGATTGACAGCTTTTTTAAACC
CAACCCATGACATGTCGAGGAAAGGGTCGTTTCGGGGAGTTAAATATTTTTGGCTATGTAGCAGACATGTTTCGACGCTGGC
GTCGCGTCGATCGGAAAATATTACCCCAGGAACAAGCACTTGCTTGGGTTAGCCACCACCCTGCGCAAGCCTTTTTGCCGGC
TCTACACAGGGCCAATGAAATCTGGGCGGAATCTGAAACCGATGAAACGGACGACACTGGCAACAAGCTCACTGCACTATT
TTATTTTTCTAGTGAAATAGCCTATCCTCGTCTCGCTCCCCTCATACCTGTAAAGGGGTGCAATTTAGCCTCGTTCCAGCCAT
TCACGGGCCACTCAACAACACGTCGGCTACCATGGGGTGCTTGGGCACCAAAAGGCCTATAAATAGGCCCCCATCCGTCTG
CTACACAGTCATCTCTGTCTTTTCTTCCCATCATACAAAATG

***$P_{HpFMD}$***

AACGAATGTATCTAAACGCAAACTCCGAGCTGGAAAAATGTTACCGGCGATGCGCGGACAATTTAGAGGCGGCGAATCAA
GAAACACCTGCTGGGCGAGCAGTCTGGAGCACAGTCATCGATGGGCCCGAGATCCCACCGCGTTCCTGGGTACCGGGACGT
GAGGCAGCGCGACATCCATCAAATATACCAGGCGCCAACCGAGTCTCTCGGAAAACAGCTTCTGGATATCTTCCGCTGGCG
GCGCAACGACGAATAATAGTCCCTGGAGGTGACGGAATATATATGTGTGGAGGGTAAATCTGACAGGGTGTAGCAAAGGT
AATATTTTCCTAAAACATGCAATCGGCTGCCCCGCGACGGGAAAAAGAATGACTTTGGCACTCTTCACCAGAGTGGGGTGT
CCCGCTCGTGTGTGCAAATAGGCTCCCACTGGTCACCCCGGATTTTGCAGAAAAATAGCAAGTTCCGGGGTGTCTCACTGGT
GTCCGCCAATAAGAGGAGCCGGCAGGCACGGAGTCTACATCAAGCTGTCTCCGATACACTCGACTACCATCCGGGTCTGTC
AGAGGGGGGAATGGCACTATAAATACCGCCTCCTTGCGCTCTCTGCCTTCATCAATCAAATCATCATACAAAATG

**Supplemental Table 3 List and sequence information of utilized signal peptides**

| *Naturally secreted protein* | *Species* | *Abbreviation* | *Sequence* |
|---|---|---|---|
| Mating pheromone alpha factor | *Saccharomyces cerevisiae* | *Sce*−Prepro | MRFPSIFTAVLFAASSALAAPVNTTTEDETAQIPAEAVIGYLD LEGDFDVAVLPFSNSTNNGLLFINTTIASIAAKEEGVSLDKREA |
| Inulinase | *Kluyveromyces marxianus* | *Kma*−Inulinase | MKLAYSLLLPLAGVSAA |
| Invertase 2 | *Saccharomyces cerevisiae* | *Sce*−Invertase2 | MLLQAFLFLLAGFAAKISA |
| Acid Phosphatase 5 | *Saccharomyces cerevisiae* | *Sce*−Acid Phosphatase | MFKSVVYSILAASLANAA |
| *Gma*UPO | *Galerina marginata* | *Gma*−UPO | MRGTPIFASLIALFAHAAIAFPAYGSLAGLTREQLDEILPTLEIRA |
| Serum Albumin | *Homo sapiens* | *Hsa*−Serum Albumin | MKWVTFISLLFLFSSAYSA |
| Glucoamylase | *Aspergillus awamori* | *Aaw*−Glucoamylase | MSFRSLLALSGLVCSGLA |
| Killer Protein K1 Toxin | *Saccharomyces cerevisiae* | *Sce*−Killer Protein | MTKPTQVLVRSVSILFFITLLHLVVA |
| *Mro*UPO | *Marasmius rotula* | *Mro*−UPO | MKLAISSSLIALVSVTTALANSQDVVDFGA |
| *Cfo*CPO | *Caldariomyces fumago* | *Cfo*−CPO | MFSKVLPFVGAVAALPHSVRA |
| *Cgl*UPO | *Chaetomium globosum* | *Cgl*−UPO | MRTSLLPALAAVSPVLA |
| *Cci*UPO | *Coprinopsis cinerea* | *Cci*−UPO | MISTSKHLFVLLPLFLVSHLSLVLGFPAYASLGGLTERQVEEYTS KLPIVA |
| α Amylase | *Aspergillus niger* | *Ani*−α Amylase | MVAWWSLFLYGLQVAAPALA |
| α Galactosidase | *Saccharomyces cerevisiae* | *Sce*−α Galactosidase | MFAFYFLTACISLKGVFGA |
| Lysozyme g2 | *Gallus gallus* | *Gga*−Lysozym | MLGKNDPMCLVLVLLGLTALLGICQGA |
| *Aae*UPO* (variant PaDa-I) | *Agrocybe aegerita* | *Aae*−UPO* | MKYFPLFPTLVYAVGVVAFPDYASLAGLSQQELDAIIPTLEARA |
| *Mth*UPO | *Myceliophthora thermophila* | *Mth*−UPO | MRASVLPVLIAISPALA |

**Supplemental Table 4 Protein sequences of the utilized UPOs and other enzymes**

**A/S-** Connecting amino acid for Golden Gate assembly

***Mhi*UPO**

UPO derived from Myceliophthora hinnulea

**A**GFDTWSPPGPTDVRAPCPMLNTLANHGFLPHDGKDITREQTENALFDALNINKTLASFLFDFALTTNPKNTSTFSLNDLGNHNIL
EHDASLSRADAYFGNVLQFNQTVFDETKTYWEGDTIDLRMAAKARLGRIKTSQATNPTYSMSELGDAFTYGESAAYVVVLGDL
ESRTVNRSWVEWFFEHEQLPQHLGWKRPAVSFEEEDLNRFMEEIEKYTKGLEGSNSTSGSQKHRRRLPRRRTHFGF**S**

***Mfe*UPO**

UPO derived from Myceliophthora fergusii

**A**GFDTWSPPGPYDVRAPCPMLNTLANHGFLPHDGKDITRENTENALFEALHINKTLGSFLFDFALTTNPRNTSTFSLNDLGNHNIL
EHDASLSRADAYFGNVLQFNQTVFDETKTYWDGDVIDLRMAARARLGRIKTSQATNPTYSMSELGDAFTYGESAAYVVVLGDK
ESRTAKRSWVEWFFEHEQLPQHLGWKRPASSLEEEDLFTIMDEIRQYTSELEGSTSSSDAQTSRRQLPRRRTHFGF**S**

***Dca*UPO**

UPO derived from Daldinia caldariorum

**A**GFAPWKAPGPDDVRGPCPMLNTLANHGFLPHDGKNIDVNTTVNALSSALNLDDELSRDLHTFAVTTNPQPNATWFSLNHLSRH
NVLEHDASLSRQDAYFGPPDVFNAAVFNETKAYWTGDIINFQMAANALTARLMTSNLTNPEFSMSQLGRGFGLGETVAYVTILG
SKETRTVPKAFVEYLFENERLPYELGFKKMKSALTEDELTTMMGEIYSLQHLPESFTKPFAKRSEAPFEKRAEKRCPFH**S**

**CalB**

lipase B derived from *Candida antarctica*

**A**LPSGSDPAFSQPKSVLDAGLTCQGASPSSVSKPILLVPGTGTTGPQSFDSNWIPLSTQLGYTPCWISPPPFMLNDTQVNTEYMVN
AITALYAGSGNNKLPVLTWSQGGLVAQWGLTFFPSIRSKVDRLMAFAPDYKGTVLAGPLDALAVSAPSVWQQTTGSALTTALR
NAGGLTQIVPTTNLYSATDEIVQPQVSNSPLDSSYLFNGKNVQAQAVCGPLFVIDHAGSLTSQFSYVVGRSALRSTTGQARSADY
GITDCNPLPANDLTPEQKVAAAALLAPAAAAIVAGPKQNCEPDLMPYARPFAVGKRTCSGIVTP**S**

**Mrl2**

laccase derived from *Moniliophthora roreri*

**A**SIGPIADLVISNQDVSPDGFTRSAVVAGGDTIGPLIVGNKNDNLQINVVNNLDDDTMLQSTSIHWHGFFQQSTNWADGTAFVNQ
CPIAKGNSFLYDFDATDQAGTFWYHSHLSTQYCDGLRGPIVIYDPDDPHASLYDVDDESTVITLADWYHTKAKEITFGTPDSTLIN
GLGRWSQGNETDLSVITVTSGQRYRMRLINTACDAAYTFSIDNHTMTVIEADAVNIEPIEVDSLTIYAGQRYSFVLNADQAVGNY
WIRANPNIGTMGYTNGINSAILRYDTAEEEEPDVLDITSTNSLSEADLVPLENPGAPGDPVVGGVDYALHLDFAFTSAATFTVNDA
TFVPPTVPVLLQILSGAQTADTLLPSGSVVALPSNSTIELSMTGGLLGLEHPIHLHGHNFDVVRVAGSTEYNYENPIRRDVVNAGS
TSDNVTIRFTTDNPGPWILHCHIDWHLEAGFAIVFAEATDEWVDTIDPSDAWENLCPTYDALSDDDLG**S**

**Supplemental Table 5 Distribution of promoters and signal peptides amongst the sequenced hits**

Overview of the occurrence and distribution of respective promoter and signal peptides parts among the sequenced top hits (80 in total) for the eight tested enzymes. Redundant combinations have not been included.

**Promoter**

| Promoter | Occurrence | Percentage |
|---|---|---|
| $P_{PpPMP20}$ | 3 | 3.8 % |
| $P_{HpMOX}$ | 4 | 5 % |
| $P_{PpFLD1}$ | 19 | 23.8 % |
| $P_{PpFDH1}$ | 11 | 13.8 % |
| $P_{PpDAS1}$ | 1 | 1.3 % |
| $P_{PpDAS2}$ | 1 | 1.3 % |
| $P_{PpCAT1}$ | 9 | 11.3 % |
| $P_{PpAOX1}$ | 2 | 2.5 % |
| $P_{PpALD4}$ | 12 | 15 % |
| $P_{HpDHAS}$ | 5 | 6.3 % |
| $P_{HpFMD}$ | 13 | 16.3 % |

**Signal Peptides**

| Signal Peptide | Occurrence | Percentage |
|---|---|---|
| *Sce*–Prepro | 3 | 3.8 % |
| *Kma*–Inulinase | 4 | 5 % |
| *Sce*–Invertase2 | 6 | 7.5 % |
| *Sce*–Acid Phosphatase | 1 | 1.3 % |
| *Gma*–UPO | 9 | 11.3 % |
| *Hsa*–Serum Albumin | 3 | 3.8 % |
| *Aaw*–Glucoamylase | 5 | 6.3 % |
| *Sce*–Killer Protein | 2 | 2.5 % |
| *Mro*–UPO | 9 | 11.3 % |
| *Cfo*–CPO | 2 | 2.5 % |
| *Cgl*–UPO | 1 | 1.3 % |
| *Cci*–UPO | 0 | 0 % |
| *Ani*–α Amylase | 8 | 10 % |
| *Sce*–α Galactosidase | 10 | 12.5 % |
| *Gga*–Lysozym | 10 | 12.5 % |
| *Aae*–UPO* | 2 | 2.5 % |
| *Mth*–UPO | 5 | 6.3 % |
